# Hidden dynamics of ubiquitin-mediated autoregulation of human 26S proteasome during substrate degradation

**DOI:** 10.1101/2020.12.22.423932

**Authors:** Zhaolong Wu, Enbo Chen, Shitao Zou, Yanzhen Hou, Shuwen Zhang, Wei Li Wang, Yuanchen Dong, Youdong Mao

**Author notes:** State Key Laboratory of Reproductive Medicine and Offspring Health, Nanjing Medical University, Nanjing, China; Department of Radiation Oncology, Nanjing Medical University Affiliated Suzhou Hospital, Gusu School, Suzhou, China (Shitao Zou). These authors contributed equally (Zhaolong Wu, Enbo Chen).

## Abstract

The 26S proteasome regulates proteostasis and myriad cellular processes. Here we reconstitute the hidden dynamics of human 26S proteasome during protein degradation using cryo-EM data-driven deep learning, which identified 64 proteasomal conformers. These conformers revealed substrate-dependent conformational entanglement of two 19S regulatory particles in the doubly capped holocomplexes. The rate-limiting step in single-nucleotide exchange dynamics of the AAA-ATPase motor is ATPase re-engagement with the substrate, while the rate-limiting step in proteasomal degradation is the initiation of substrate unfolding by the AAA-ATPase motor following deubiquitylation. Nine rare conformers delineated novel ubiquitin-binding sites on the RPN2 and α5 subunits, along with a ubiquitylation site on RPN10, which were verified using site-directed mutagenesis and liquid chromatography-tandem mass spectrometry. Opening of the 20S core particle gate was found to upregulate proteolytic activity by allosterically promoting nucleophilic attack on the scissile peptide bond. Our systematic analyses illuminate a grand hierarchical allostery underlying ubiquitin-mediated proteasome autoregulation.

## Introduction

The 26S proteasome is the known largest ATP-dependent protease machinery of ∼2.5 MDa molecular weight ubiquitously found in all eukaryotic cells. It exists at the center of the ubiquitin-proteasome system (UPS) that regulates myriad cellular processes, such as protein quality control, cell cycle, gene expression, inflammation and immunity.^1–3^ Dysregulation of the UPS has been broadly associated with carcinogenesis, ageing, neurodegenerative and cardiovascular diseases.^2–5^ The 26S holoenzyme is assembled by a cylindrical 20S core particle (CP) capped with one or two 19S regulatory particles (RPs), each consisting of the lid and base subcomplexes.^2,6^ The ring-like heterohexameric motor of ATPases-associated-with-diverse-cellular-activities (AAA) ATPase in the base subcomplex mechanically unfold polyubiquitylated substrates recognized by ubiquitin receptors in the lid subcomplex.

Previous structural studies by cryogenic electron microscopy (cryo-EM) have mapped the major conformational states of substrate-engaged proteasome.^6–8^ Several modes of coordinated ATP hydrolysis in the AAA-ATPase motor were observed to regulate distinct steps of ubiquitin recognition (states E_A1_ and E_A1_), deubiquitylation (state E_B_), initiation of translocation (states E_C1_ and E_C2_) and processive substrate translocation (states E_D1_ and E_D2_) in the proteasome.^6–8^ The incorporation of the deubiquitylating enzyme USP14 into the proteasome reprograms this pathway by overwriting states E_B_, E_C1_ and E_C2_, and stabilizes translocation- competent E_D_-like states^8^. However, key intermediate conformations connecting major states were largely missing, limiting our understanding of the intrinsic proteasome regulation. Moreover, the mechanism of the proteasome targeting of substrate-conjugated ubiquitin signals is incompletely understood.^2,9–11^ How polyubiquitylated substrate interactions regulate proteasome activity remains enigmatic.

The 26S proteasome is one of the most complex, dynamic, conformationally and compositionally heterogeneous holoenzyme machineries in cells^2,6,7,12–14^, challenging the current approaches in cryo-EM structure determination.^9,15,16^ Visualizing atomic structures of transient, nonequilibrium intermediates connecting the major metastable states of the proteasome has been unsuccessful with conventional cryo-EM analysis due to extreme conformational heterogeneity.^2,6,7^ To address this major challenge, we recently developed a deep manifold learning framework named AlphaCryo4D,^17^ which significantly improved the accuracy of 3D classification. This method breaks existing limitations of resolution in reconstructing continuous conformational motions and enables atomic-level cryo-EM reconstructions of highly dynamic, lowly populated conformational intermediates or transient states. Recent application of this approach in a time-resolved cryo-EM study has comprehensively revealed detailed molecular mechanism of the mutual regulation between the deubiquitylating enzyme USP14 and the human 26S proteasome that were further substantiated by functional and mutagenesis experiments.^8^

In this Article, we report 64 conformers of the human 26S proteasome in the act of substrate degradation by using cryo-EM data-driven deep learning in combination with complementary experimental and computational approaches such as site-directed mutagenesis, microscale thermophoresis (MST), liquid chromatography-tandem mass spectrometry (LC–MS/MS), molecular dynamics simulation and quantum mechanical calculation. Systematic analyses of these proteasomal conformers in substrate processing pinpoint energetic differences between the doubly and singly capped proteasomes, discover several novel ubiquitin-binding sites in the proteasome, and choreograph single-nucleotide-exchange dynamics of the proteasomal AAA-ATPase motor during translocation initiation. Importantly, these enriched proteasomal states provide unprecedented insights into an integrative, hierarchical allosteric mechanism of proteasome autoregulation upon polyubiquitylated substrate interactions.

## Results

### Visualizing hidden dynamics of the proteasome

We performed deep-learning-based 3D classification on a large cryo-EM dataset of the substrate-engaged 26S proteasome using AlphaCryo4D^17^ (see Methods; Table S1). This dataset was collected from the human 26S proteasome mixed with a Lys63-linked polyubiquitinated substrate Sic1^PY^ for 30 seconds before ATP was exchanged to ATPγS, which was expected to maximally capture any intermediate states before the degradation reaction was completed.^6^ Initial analysis of this dataset using conventional approaches in cryo-EM data analysis has previously yielded seven atomic structures of the proteasome^6^. The energy landscape of the substrate-engaged proteasome reconstituted by our AlphaCryo4D procedure using all available single-particle data (see Methods for details) exhibits the highest activation energy barrier between states E_B_ and E_C1_ (Figures 1B and 1C), suggesting that the rate-limiting step of substrate degradation lies in the initiation of substrate unfolding after substrate deubiquitylation, consistent with recent fluorescence microscopy studies^18^. Comparison with the energy landscape of the substrate-free ATPγS-bound 26S proteasome^14^ indicates that substrate interactions considerably lower the relative free energy of the substrate-engaged E_D_-like states, thus shifting the conformational space of the proteasome toward the states of open CP gates (Figure 1A).

**Figure 1.**
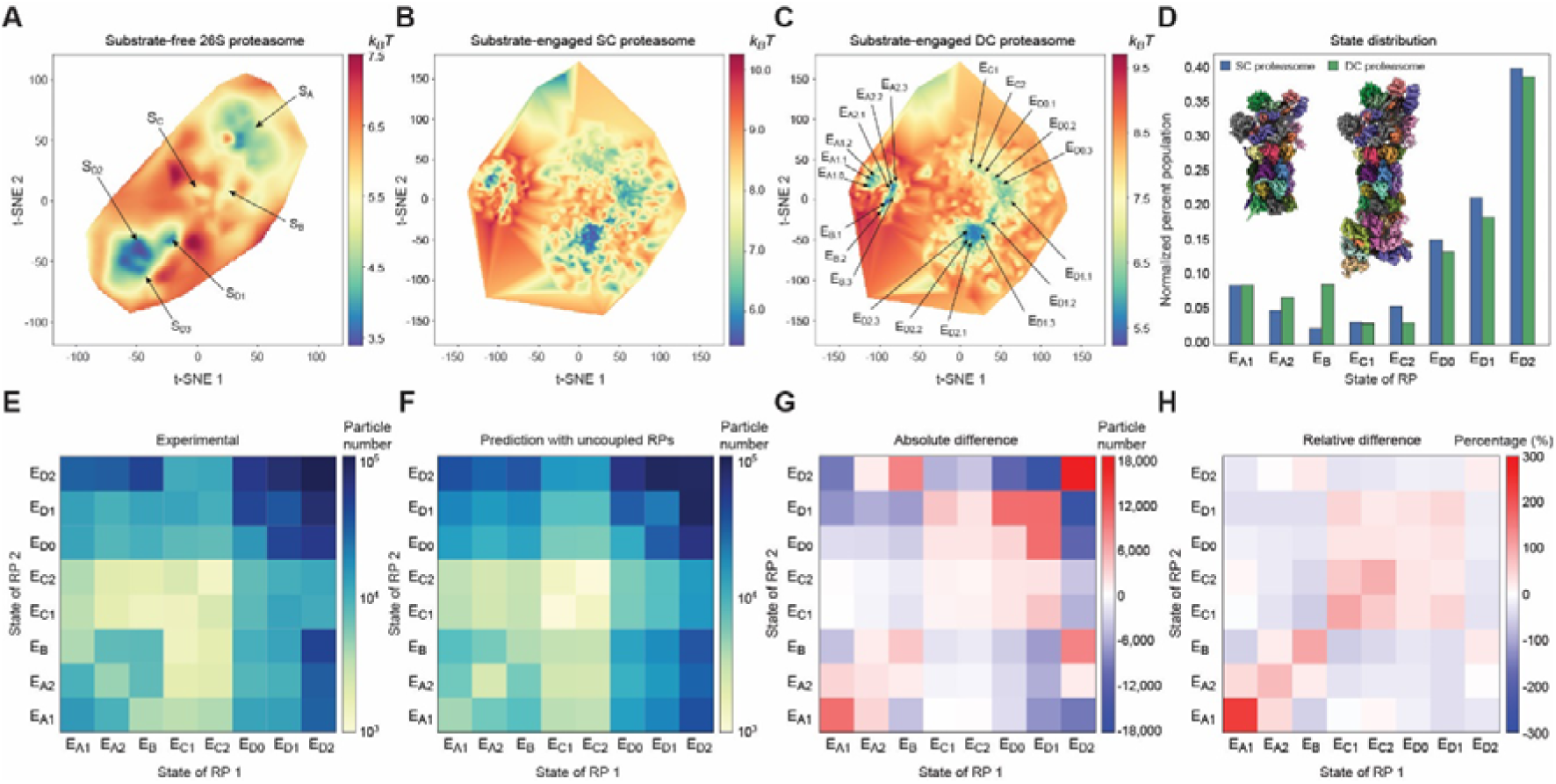
Energetic differences of the singly and doubly capped human proteasomes. (**A**-**C**) Energy landscape of the substrate-free ATPγS-bound 26S proteasome (**A**), the substrate-engaged SC proteasome (**B**), and the RP-CP subcomplex of the substrate-engaged DC proteasome (**C**). The energy landscape of the substrate-free proteasome shows six local energy minima corresponding to the known six conformational states S_A_, S_B_, S_C_, S_D1_, S_D2_ and S_D3_.^14^ In contrast, 20 conformational states of the RP-CP subcomplex are marked on the energy landscape of the substrate-engaged 26S proteasome. (**D**) Comparison of the state distribution of RP in the SC and DC proteasomes. (**E**) State distribution matrix of the DC proteasome obtained using experimental data, colored by the particle numbers in the DC states, with the horizonal and vertical axes representing the states of two RPs bound to the same CP. (**F**) State distribution matrix of the DC proteasome predicted by a control model assuming the two completely uncoupled RPs in the same DC proteasome. The model assumes the same total number of DC particles and the same probability of observing each RP state as experimentally measured. (**G**) State distribution difference matrix of the DC proteasome by experimental results minus the model predictions, colored by the absolute particle number difference. (**H**) Relative differences of the DC proteasome computed as the ratio of absolute difference divided by the particle numbers from the model prediction in each DC state.

Our deeper 3D classification allowed reconstructions of significantly more conformers of the proteasomes with numerous novel features compared to the previously reported results using traditional methods.^6^ It discovered 20 conformational states of RP-CP subcomplex (Figures 1B, 1C, 2 and S2; and Table S1). Nine lowly populated states (designated E_A1.1_, E_A1.2_, E_A2.2_, E_A2.3_, E_B.2_, E_B.3_, E_D2.2_, E_D2.3_ and E_D.α5_) at 3.1-7.5 Å resolution exhibit previously unseen features resembling ubiquitin that map various putative ubiquitin-binding sites on the RPN1, RPN2 and α5 subunits as well as putative ubiquitylation of RPN10 in the proteasome (Figures 2-4, S1F–S1K, and Movie S1). Six sequential intermediate states (designated E_D0.1_, E_D0.2_, E_D0.3_, E_D1.1_, E_D1.2_ and E_D1.3_) at 3.2-3.9 Å resolution bridge a major missing gap between states E_C2_ and E_D1_, revealing how concerted structural transitions in the AAA-ATPase motor during single-nucleotide exchange drive the initiation of substrate unfolding (Figure 5, Movie S2).

**Figure 2.**
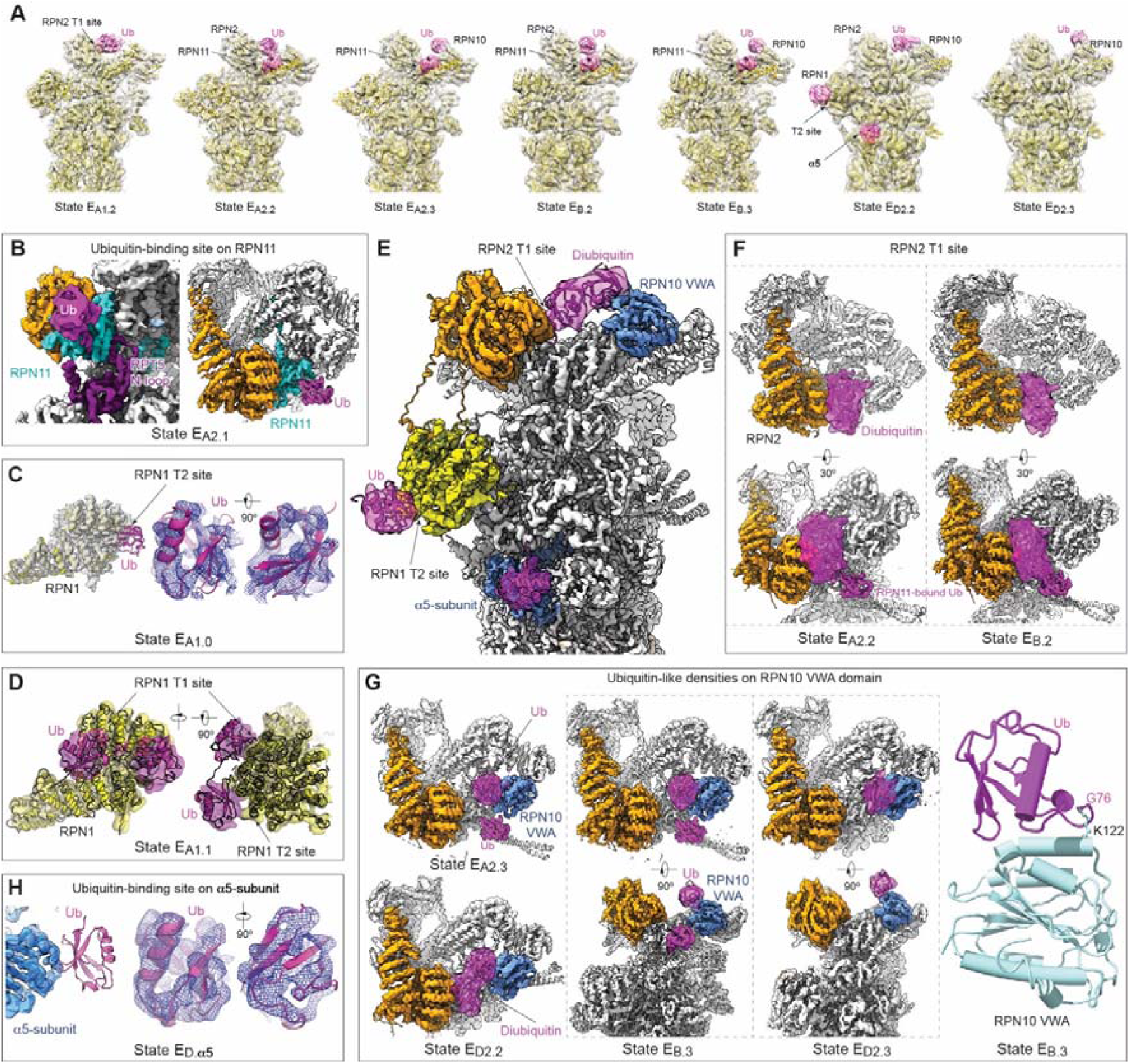
Cryo-EM visualization of ubiquitin-proteasome interactions in various conformational states captured during substrate degradation. (**A**) Seven lowly populated states visualize various ubiquitin-binding sites on the RPN1, RPN2, RPN10 and α5 subunits. For clarity, the cryo-EM maps are low-pass filtered at 8 Å and shown in transparent surface representations superimposed with their corresponding atomic models in cartoon representations. The ubiquitin (Ub) densities are colored magenta. (**B**) Cryo-EM density of state E_A2.1_ shows the improved resolution localized around RPN11-bound ubiquitin and the RPT5 N-loop as compared to previously determined state E_A2_. Ub, ubiquitin. (**C**) The refined 3.7-Å RPN1 density map from state E_A1.0_ in transparent surface representation superimposed with its atomic model with Ub bound near the RPN1 T2 site. The right insets show the local cryo-EM density of the RPN1-bound Ub in state E_A1.0_ superimposed with its atomic model in cartoon representation. (**D**) Cryo-EM density of state E_A1.1_ in transparent surface representation superimposed with its pseudo-atomic model of cartoon representation, with one Ub moiety bound to the RPN1 T1 site and another Ub near the T2 site. The two Ub moieties appear to be compatible with a Lys63-linked diubiquitin model in an extended conformation, which is supported by the low-level density connecting the two Ub densities. (**E**) High-resolution cryo-EM density of state E_D2_ superimposed with Ub densities in state E_D2.2_ highlighted as magenta, showing that a diubiquitin chain in contact with both the RPN2 T1 site and the RPN10 VWA domain. The atomic model of state E_D2.2_ in cartoon representation is superimposed with the density map. (**F**) Closeup view of the RPN2-bound diubiquitin density in states E_A2.2_ (left column) and E_B.2_ (right column) in two different perspectives, with the lower row showing the coexistence of the RPN11-bound Ub and the RPN2-bound diubiquitin. (**G**) Closeup views of the RPN10-associated Ub densities in states E_A2.3_, E_B.3_, and E_D2.3_. Right panel shows the pseudo-atomic model of ubiquitylation at Lys122 in the RPN10 VWA domain. (**H**) Closeup view of the α5-subunit bound with Ub in the refined 3.1-Å density map of state E_D.α5_. The cryo-EM density of the α5-subunit is shown in transparent surface representation superimposed with its atomic model. The right insets show the local cryo-EM density of the α5-bound Ub of state E_D.α5_ in blue mesh representation low-pass filtered at 4.9 Å and superimposed with its atomic model in cartoon representation.

To understand whether the singly capped (SC) and doubly capped (DC) proteasomes behave differently, we reconstructed 8 and 36 conformational states for the pure SC and DC holoenzymes, respectively (Figures 1D, 1E, S3 and S4). All 8 SC and 29 DC states were refined to 3.7-4.7 Å, with 7 DC states limited to 6-9 Å due to their extremely low population (Figures S3B, S3E, and S3F). Note that the CP gate in the RP-distal side of the 20 states of RP-CP subcomplex is in a heterogeneous conformation of various stoichiometric ratios between closed and open states. By contrast, both CP gates in the 44 states of pure SC or DC reconstructions are in a homogeneous conformation.

Besides resolving significantly more conformers, improved 3D classification pushes the envelope of the achievable resolution of known states by keeping more particles without sacrificing conformational homogeneity. For example, the local resolution of RPN11-bound ubiquitin in state E_A2.1_ was improved from 5-6 Å to 3.8 Å (Figure 2B). It reveals a previously unsolved feature of a short substrate stretch bound to the cleft between RPN11 and RPT4 at the ATPase entrance prior to substrate insertion into the ATPase ring in state E_B_ (Figure S5). The structure of state E_D2_ was also improved from the previous resolution at 3.2 Å to 2.5 Å, allowing us to investigate how the CP gating allosterically regulates the catalytic activity of the β-type subunits (Figure 6).

### Conformational entanglement

Both the SC and DC proteasomes are abundant in cells.^19^ Their structural and functional difference, however, remains elusive. In the 36 DC reconstructions, each conformer is a combination of two RP states. Each CP gate in the DC proteasome appears to be rigorously controlled by their respective proximal RP and is only open when its proximal RP is in an E_D_-compatible state with five RPT C-terminal tails inserted into the surface pockets on the α-ring (Figures 1E and S3A-S3D).^6^ Consistently, the gate on the RP-free side of the CP in the SC proteasome is closed in all states, indicating that the CP gate opening on the RP-controlled side does not remotely open the other CP gate on the RP-free side (Figure S3D).

Previous studies have found no obvious coupling of two RP conformations in the same DC proteasome in the absence of substrate.^14,19^ To understand whether this is also true in the presence of substrate, we compare the state distribution matrix obtained by experimental data to the prediction of a control model assuming two completely uncoupled RP states in the DC proteasome (Figures 1F-1H). Although the overall pattern of the experimental distribution roughly agrees with the model prediction, a notable number of DC states exhibit substantial deviation, indicating that the conformational dynamics of the two RPs in the same proteasome are profoundly entangled together (Figures 1G and 1H). Intriguingly, the most prominent positive deviations are seen in the state distributions of E_A1_-E_A1_, E_B_-E_B_, E_B_-E_D2_, and E_D2_-E_D2_ combinations (Figure 1G), suggesting that the deubiquitylation state E_B_ and the translocation state E_D2_ of one RP are allosterically coupled to and mutually upregulated by states E_B_ and E_D2_ of the opposite RP. The state distribution of DC proteasome with E_A1_-E_A1_ configuration appears to be disproportionately enriched, presumably due to lack of substrate in the AAA-ATPase ring. Coupled actions of substrate deubiquitylation and translocation between two opposite RPs are expected to optimize the efficiency of substrate processing. Moreover, all the DC states along the diagonal of the state distribution matrix are upregulated (indicated by red in Figures 1G and 1H), suggesting that both RPs in the same proteasome synergistically promote each other to transit into downstream states in a symmetrical fashion.

Notably, the largest discrepancy of the distribution of RP states between the SC and DC proteasomes also lies in state E_B_ (Figures 1B-1D). The state population of E_B_ is significantly lower in the SC than in the DC proteasome. This observation verifies the conclusion drawn from the above analysis on the DC proteasomes alone that state E_B_ is upregulated by the conformational entanglement effect in the DC holoenzyme, suggesting that the SC proteasome lacks the functional synergy between two opposite RPs seen in the DC proteasome (Figure 1H). Altogether, these results implicate that the DC proteasome could be energetically more efficient in substrate deubiquitylation and degradation than the SC proteasome.

### Ubiquitin-RPN1 interactions

By focused 3D classification, we refined the density map of RPN1 in state E_A1.0_ (∼5.9% of the dataset) to 3.7 Å, improving upon the previous reconstruction at moderate resolution (∼5.5 Å) (Figures 2C and S1D).^6^ This map reveals that a putative ubiquitin moiety contacts Glu458, Cys459 and Asp460 at the N-terminal end of an RPN1 helix (homologous to H24 between the T1 and T2 sites of yeast Rpn1) and potentially interacts with the T2 site (Asp423, Leu426 and Asp430) on the RPN1 toroidal domain (Figures 2C and S1F).^11^ Since the yeast Rpn1 T2 site was found to preferentially bind Ubp6 ubiquitin-like (UBL) domain^11^, this site may be sterically occluded for ubiquitin binding in the presence of USP14/Ubp6. In addition, we also identified another state E_A1.1_ that exhibits a diubiquitin-like density with one ubiquitin moiety on the RPN1 T1 site and the other near the T2 site (Figure 2D). The much smaller population of state E_A1.1_ (∼1.3 % of the dataset) is consistent with the previous finding that Lys63-linked polyubiquitin chains have a lower affinity against the T1 site than those linked via Lys48 and Lys6 residues.^11^

### Ubiquitin-RPN2 interactions

Unexpectedly, in states E_A1.2_, E_A2.2_, and E_B.2_, a diubiquitin-like density is consistently found on RPN2, whereas a monoubiquitin-like density is observed at the apex of the von Willebrand factor type A (VWA) domain^20^ of RPN10 in states E_A2.3_, E_B.3_ and E_D2.3_ (Figures 2A). In state E_D2.2_, the diubiquitin-like density, however, appears to contact both RPN2 and RPN10, and can be approximately fitted with a diubiquitin (Figure 2E and 2F). The RPN10 ubiquitin-interacting motifs (UIMs), however, were not reliably observed presumably due to their highly flexible nature (Figure 2G). The populations of these states are very small and represented in only a few thousand particles or 0.1-0.3% of the entire dataset, indicating their transient nature. The local resolutions of the RPN2- and RPN10-bound ubiquitin-like densities are lower than the rest of the reconstructions and other ubiquitin densities on RPN1, RPN11 and α5 subunits. Rigid-body fitting of the ubiquitin structure into these ubiquitin-like densities confirms the agreement of the overall density shape with the ubiquitin fold (Figure 2).

The ubiquitin-binding site on RPN2 is centered on three helices harboring an acidic surface around residues Tyr502, Glu530, Asp531 and Ser569 (Figures 3A-3D). Structural comparisons suggest that this site on RPN2 is homologous to the T1 site of RPN1 (Figures 3B and 3C)^11^. We therefore refer to it as ‘the RPN2 T1 site’. Although there is little conservation in amino acid sequences between the RPN1 and RPN2 T1 sites, their structural and electrostatic properties are highly comparable (Figures 3C, S6A–S6D). Comparison with the NMR structure of Lys48-linked diubiquitin-bound RPN1 T1 site helices^11^ further confirms the similarity of the ubiquitin-binding modes between the RPN1 and RPN2 T1 sites (Figure 3B). Consistent with this finding, previous studies have observed weak interactions between UBL proteins and RPN2 via cross-linking experiments.^21^

**Figure 3.**
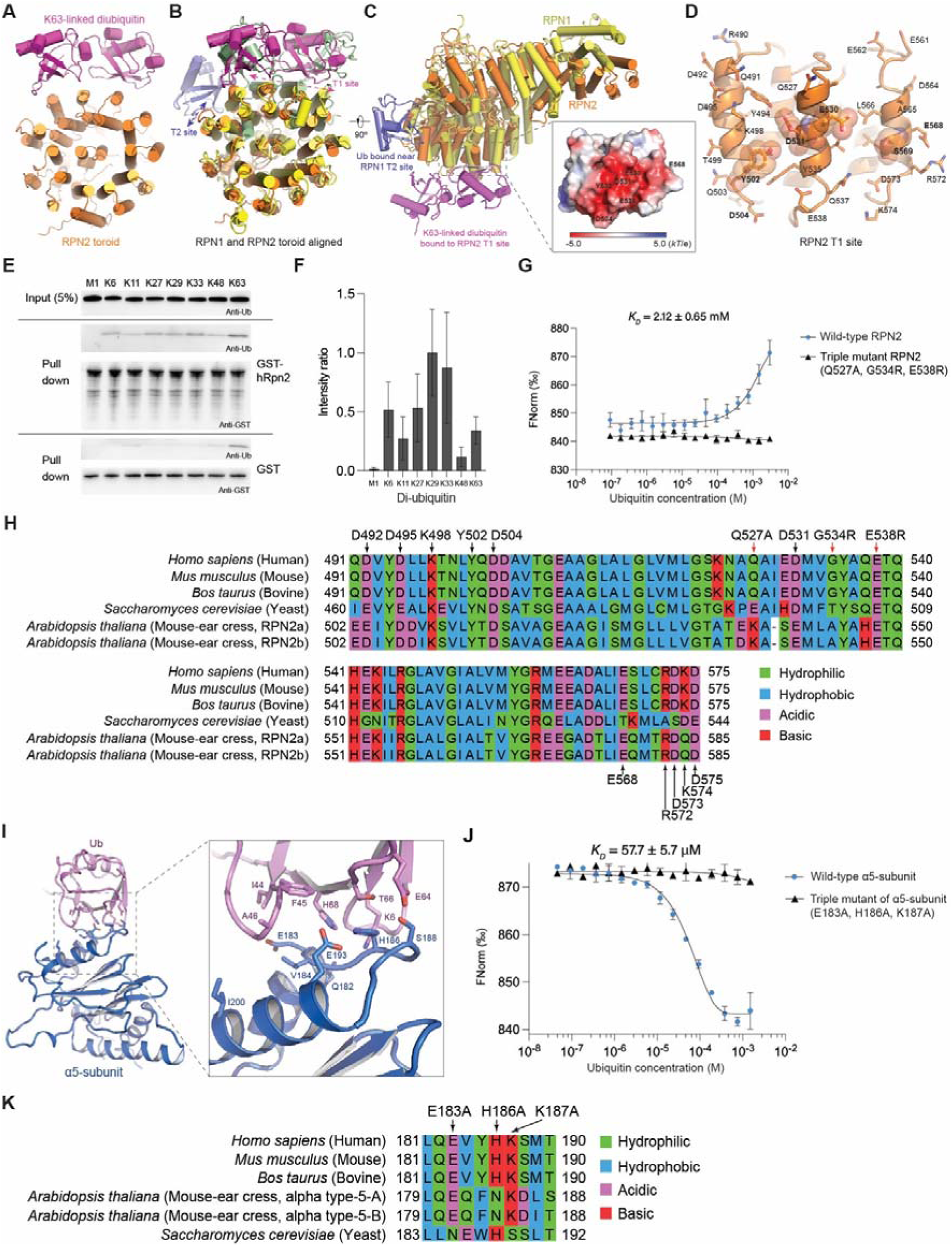
Ubiquitin-binding sites on RPN2 and α5 subunits. (**A**) Structure of the RPN2 toroidal domain (orange) in complex with Lys63-linked diubiquitin (magenta) modelled from the cryo-EM maps of states E_A1.2_, E_A2.2_ and E_B.2_, in cartoon representation. (**B**) Structure-based alignment of the human RPN1 (yellow) and RPN2 (orange) toroidal domains. Ubiquitin bound near the RPN1 T2 site is shown as light blue. The NMR structure (PDB ID 2N3V) of the Lys48-linked dibuiquitin-bound yeast Rpn1 T1 site segment,^11^ shown in light green, is aligned and superimposed with the human RPN1 and RPN2. (**C**) The aligned human RPN1 and RPN2 are viewed from a perspective orthogonal to that in panel (**B**). Insert, electrostatic surface of the RPN2 T1 site showing its acidic nature. (**D**) Closeup view of the RPN2 T1 site composed of three helices in carton representation with the side chains shown in stick representation. The key residues contacting ubiquitin are highlighted with transparent sphere representation. (**E**) Pull-down assay with full-length GST-RPN2 or GST (negative control) protein and M1-, K6-, K11-, K27-, K29-, K33-, K48-, and K63-linked diubiquitin. Immunoblotting was done with antibody against ubiquitin (anti-ubiquitin) (first, second and fourth panels) or antibody against GST (anti-GST) (third and fifth panels). Direct loading for 5% of the diubiquitin input for each chain type with immunoblotting by ubiquitin-specific antibody is included (top, left), while the results of pull-down assay with full-length GST-RPN2 (experimental group, middle, left) and GST (negative control group, bottom, left) protein are showed below the input results. (**F**) The pull-down assay was repeated four times, and the diubiquitin signal intensities were separately normalized to the strongest signal by using ImageJ. The average value and standard deviation are plotted. (**G**) MST analysis of ubiquitin binding to the human RPN2. A dissociation constant of 2.12 ± 0.65 mM for wild-type RPN2 were calculated from two independent experiments (shown as mean ± s.d.). (**H**) Homology sequence alignment of RPN2 T1 site sequence from five different species. The amino acids pointed by the arrows are the amino acids that may be involved in ubiquitin binding, while the amino acids pointed by the red arrows are the mutated amino acids. Amino acids with green background are hydrophilic amino acids, amino acids with blue background are hydrophobic amino acids, amino acids with pink background are acidic amino acids and amino acids with red background are basic amino acids. (**I**) Structure of ubiquitin-bound α5 subunit shown in cartoon representation. The side chains of residues mediating intermolecular interactions are shown in stick representation. (**J**) MST analysis of ubiquitin binding to the human α5 subunit. A dissociation constant of 57.7 ± 5.7 μM for wild-type α5 subunit were calculated from two independent experiments (shown as mean ± s.d.), whereas the triple mutant of α5 shows no obvious ubiquitin binding. (**K**) Homology sequence alignment of α5 from five different species. The range of multiple sequence alignment is shown at the beginning and end of each sequence. The amino acids pointed by the arrows are the amino acids that may be involved in ubiquitin binding, while the amino acids pointed by the red arrows are the mutated amino acids. Amino acids with green background are hydrophilic amino acids, amino acids with blue background are hydrophobic amino acids, amino acids with pink background are acidic amino acids and amino acids with red background are basic amino acids.

Ubiquitin forms polyubiquitin chains through seven lysine residues (Lys6, Lys11, Lys27, Lys29, Lys33, Lys48 and Lys63) or the N-terminal methionine (Met1). Polyubiquitin chains of different linkage types have distinct structural and signaling properties.^22^ To evaluate the preference of human RPN2 against ubiquitin-chain linkage types, resin-bound GST-RPN2 or GST (negative control) was incubated separately with diubiquitin of each linkage type, washed, and assayed for complex formation by immunoblotting with a ubiquitin-specific antibody (Figure 3E). In contrast to RPN1’s preference for Lys6- and Lys48-linked ubiquitin chains^11^, RPN2 demonstrated a preference for Lys29- and Lys33-linked diubiquitin, with the second lowest affinity for Lys48-linked diubiquitin (Figure 3F). Nonetheless, both RPN2 and RPN1 exhibit the weakest recognition for Met1-linked diubiquitin.

To quantify the interaction between RPN2 and ubiquitin, we used microscale thermophoresis (MST) to measure the dissociation constant (*K*_D_) of separately purified full-length human RPN2 and ubiquitin. Multiple sequence alignment of RPN2 from various species including yeast, plant, and human indicates a high degree of conservation of the RPN2 T1 site in eukaryotes (Figure 3H). To verify the role of the RPN2 T1 site for ubiquitin binding, we introduced mutations at three conserved residues (Q527A, G534R and E538R) within the RPN2 T1 site, and analyzed the interactions of both the triple-mutant and wild-type RPN2 with ubiquitin using MST. By fitting the MST curve of wild-type RPN2, the *K*_D_ value for the interaction between wild-type RPN2 and ubiquitin was measured to be 2.12 ± 0.65 mM. In contrast, the MST curve of triple mutant RPN2 remains flat over the measured concentration range and shows no observable binding, indicating that the mutations at the RPN2 T1 site severely disrupt ubiquitin binding. This result is further supported by all-atom molecular dynamics simulation of ubiquitin in complex with either wildtype or triple mutant RPN2 (Figure S6I). Taken together, our data suggest that the RPN2 T1 site serves as a low-affinity binding site for ubiquitin, with a linkage preference complementary to that of RPN1.

### Ubiquitin-**α**5 interaction

In state E_D2.2_, we can observe a ubiquitin density bound to the RPN1 T2 site and another weaker ubiquitin density at the α5-subunit ∼6 nm beneath the RPN1 T2 site (Figures 2A and 2E). To determine whether the ubiquitin-binding site on the α5 subunit is used more broadly, we combined all the E_D_-compatible states and focused the 3D classification on the α5-bound ubiquitin. This yielded a larger 3D class, designated as state E_D.α5_ (∼4% of the entire dataset), which improved the resolution of the ubiquitin-bound CP structure to 3.1 Å, and the α5-bound ubiquitin density to 4.9 Å (Figures S1E, S1H, S1F, S1K and S3F). The ubiquitin-binding site on the α5 subunit is composed of charged or hydrophobic residues Glu183, Val184, His186, Lys187 and Glu193 in a loop connecting two helices, resembling the mode of ubiquitin recognition by an RPN13 loop (Figures 2H and 3I).^10,23^ The electrostatic property of this site is also similar to those of the RPN1 T1/T2 and RPN2 T1 sites, which are acidic and in charge complementarity with the basic ubiquitin surface around residues Ile44 and His68 (Figures S6A–S6H). Although no previous evidence supports ubiquitin interactions with the CP, recent data suggest that the free 20S CP can degrade substrate-conjugated ubiquitin, which might be facilitated by ubiquitin binding to the α-ring.^24^

To examine the interaction between the α5 subunit and ubiquitin, we used microscale thermophoresis (MST) to measure their dissociation constant (*K*_D_). We analyzed the interactions of both wild-type and triple mutant α5-subunit (E183A, H186A and K187A) with ubiquitin. By fitting the MST curve of wild-type α5-subunit, the *K*_D_ for the interaction between wild-type α5-subunit and ubiquitin is measured to be 57.7 ± 5.7 μM. In contrast, the MST curve of the triple mutant α5 remained essentially flat over the given concentration range, indicating that the mutations abolished the ubiquitin-α5 interaction. These results confirm the ubiquitin-binding site of α5, which consists of a charged loop (Glu183-Ser188) connecting two helices (Ser174-Ser180 and Leu191-Gln204). Notably, the *K*_D_ of RPN2 and ubiquitin is two orders of magnitude larger than that of the α5 subunit. This is consistent with the much smaller population of conformers with RPN2 bound to ubiquitin.

Furthermore, we performed all-atom molecular dynamics simulation of the α5- ubiquitin system for 100 ns. Using the coordinate system of α5 as a reference, we calculated the RMSD of ubiquitin as a function of time, reflecting its overall structural motions. Notably, the RMSD of ubiquitin bound to the wild-type α5 was substantially lower than that of ubiquitin bound to triple mutant α5 (Figure S7J), suggesting that triple mutations at the three highly conserved sites (Figure 3K) disrupt ubiquitin binding to α5. These results further support the ubiquitin-binding site on α5 subunit.

### Ubiquitylation of the 26S proteasome

In states E_A2.3_, E_B.3_ and E_D2.3_, we consistently observed monoubiquitin-like densities on Lys122 of the RPN10 VWA domain. By contrast, in state E_D2.2_, a diubiquitin-like density appears to span from this lysine-rich site of RPN10 to the RPN2 T1 site and can be tentatively modelled with a Lys63-linked diubiquitin (Figures 2E–2G). To understand the nature of these additional ubiquitin-like densities, we performed liquid chromatography–tandem mass spectrometry (LC–MS/MS) analysis on the ubiquitylation of purified human 26S proteasome. Consistent with previous reports,^25,26^ the LC-MS/MS detected various proteasome-associated proteins, including several E3 ubiquitin ligases UBE3A and UBE3C. Importantly, the LC-MS/MS results revealed that several lysine residues of the RP subunits have been covalently modified by ubiquitin, including Lys292 in RPN2, Lys38 in RPN3, Lys122 in RPN10, Lys21 and Lys34 in RPN13, Lys20 in RPT4 and Lys70 in RPT5 (Figure 4). Unexpectedly, we also observed a serine ubiquitylation at Ser19 of RPN13, whose physiological function requires further investigation (Figure 4E). Although our cryo-EM analysis did not capture the ubiquitin densities resulting from ubiquitylation of the proteasomal subunits except for RPN10, pseudo-atomic modeling of Lys122- linked ubiquitin on RPN10 indeed agrees with the ubiquitin-like densities observed in the cryo-EM maps (Figure 2G).

**Figure 4.**
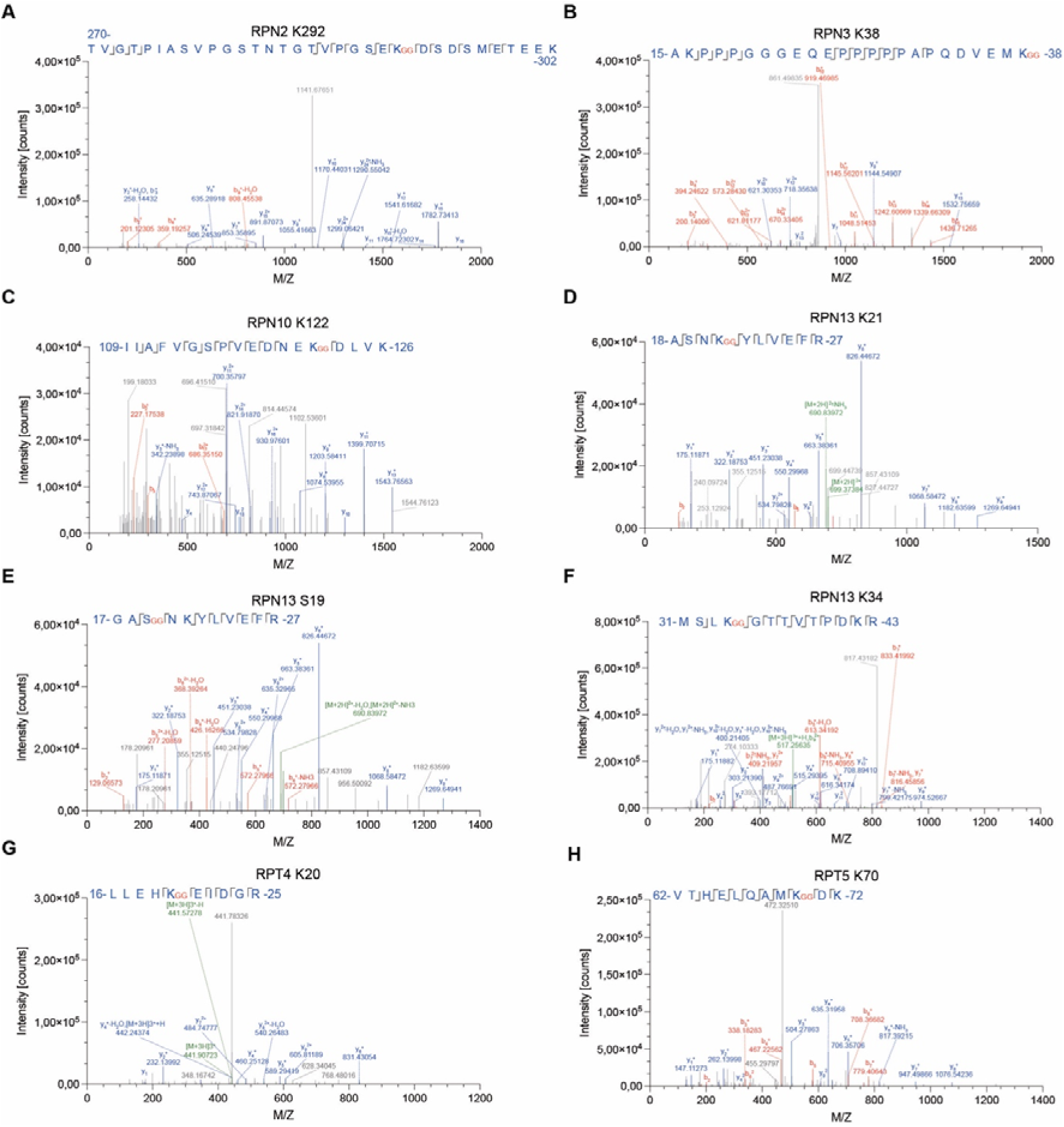
Ubiquitylation analysis of the human 26S proteasome by LC-MS/MS. Liquid chromatography–tandem mass spectrometry (LC–MS/MS) assay was used to analyze the ubiquitination of 26S proteasome subunits, and revealed that several protein subunits have ubiquitin modification sites including Lys292 in RPN2 (**A**), Lys38 in RPN3 (**B**), Lys122 in RPN10 (**C**), Lys21 (**D**), Ser19 (**E**) and Lys 34 (**F**) in RPN13, Lys20 in RPT4 (**G**) and Lys70 in RPT5 (**H**).

In line with these observations, previous studies have suggested that the yeast Rpn10 VWA is a ubiquitin-binding domain and certain ubiquitylation of Rpn10 VWA itself prevents Rpn10 from being incorporated into the proteasome.^20^ Ubiquitylation of RPN10 has been found to regulate substrate recruitment.^20,27–29^ RPN10 has also been found to mediate autophagic degradation of 26S proteasome itself.^30,31^ Ubiquitylation of the proteasome is used for the clearance of inhibited or malfunctional proteasome subunits in cells. It is possible that some sites of ubiquitylation detected in our LC-MS/MS experiments are involved in proteasomal autophagy for self-clearance. Whether the specific ubiquitylation of proteasomal subunits is functionally required beyond self-clearance deserves further investigation.

### Single-nucleotide exchange dynamics

The six intermediate states (E_D0.1_, E_D0.2_, E_D0.3_, E_D1.1_, E_D1.2_ and E_D1.3_) during initiation of substrate unfolding and translocation exhibit highly coordinated movements in the RP relative to the CP (Figures 5A–5C and S7). There are prominent rotations in the RPN1 N-terminal domain, which detaches from RPT2 in states E_D0.1_, E_D0.2_, and E_D0.3_ and re-binds RPT2 in state E_D1.1_, E_D1.2_ and E_D1.3_. All six conformations share the same architecture in their substrate interactions with pore-1 loops from four RPT subunits (RPT2, RPT6, RPT3 and RPT4), which remain largely unchanged in overall conformation (Figure 5D). These structures illustrate that the RPT2 re-association with the substrate proceeds that of RPT1 after both RPT1 and RPT2 have dissociated from the substrate in state E_C_. They also share the same pattern of nucleotide states, with ATP bound to RPT1, RPT2, RPT6 and RPT3 and ADP bound to RPT4. There are poor, partial nucleotide densities in the RPT5 nucleotide-binding pocket, which give rise to the highest B-factor (250-500) when fitted with ADP, suggesting that RPT5 undergoes ADP release in these states (Figure S7K).

**Figure 5.**
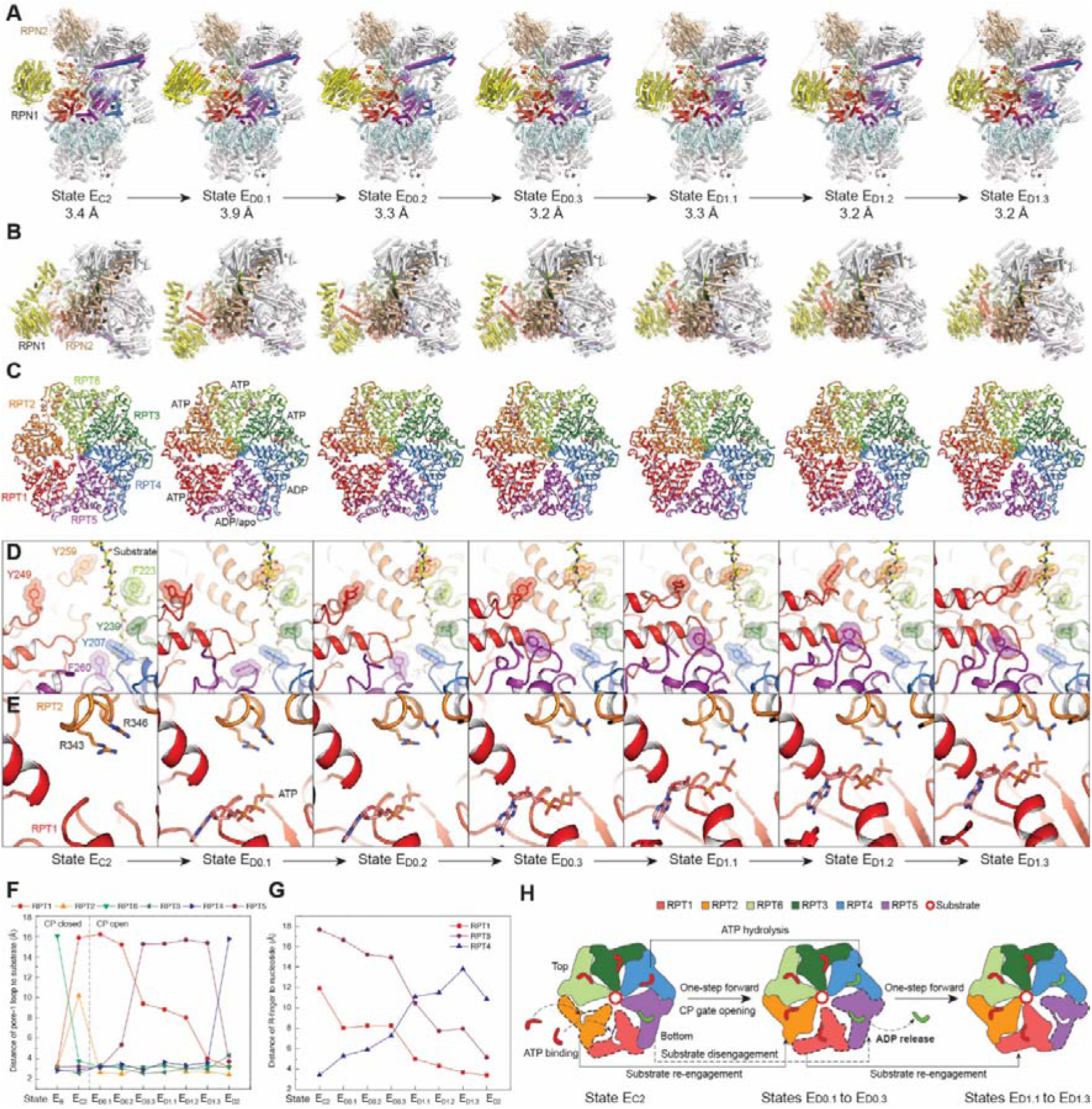
Single-nucleotide-exchange dynamics of the proteasomal AAA-ATPase motor during translocation initiation. (**A**) Side views of the substate- engaged RP-CP subcomplex of seven sequential conformers in cartoon representation. (**B**) Top views of the seven sequential conformers, showing the overall lid rotation and the relative rotation between RPN1 and the rest of the RP structure. (**C**) The AAA-ATPase structures of the seven conformers in cartoon representations. The substrates and nucleotides are shown in stick representations. (**D**) Close-up views of the pore-1 loop interactions with the substrate (in yellow stick representation), with the aromatic residues in the pore-1 loops highlighted by transparent sphere representations. (**E**) Close-up views of the RPT1 nucleotide-binding sites in different states. (**F**) Plots of distances of the pore-1 loops of all RPT subunits to the substrate measured in the atomic structures of 9 sequential conformational states. (**G**) Plots of distances of R-finger to the nucleotides bound to three RPT subunits measured in the atomic structures of 8 sequential conformational states. (**H**) Schematic illustrating the coordinated ATP hydrolysis reaction and nucleotide exchange associated with the intermediates during the transition from state E_C2_ to E_D1.3_.

Markedly, the pore-1 loop of RPT1 gradually moves toward the substrate from a distance of approximately 16 Å to 3 Å, while the pore-1 loop of RPT5 gradually moves away from the substrate from approximately 3 Å to 16 Å (Figure 5F). The small AAA subdomain of RPT5 moves inward together with the large AAA subdomain of RPT1, as the large AAA subdomain of RPT5 rotates away from the substate (Figure S7F-I). Consistently, the arginine fingers (R-finger) of RPT2 progressively move toward the ATP bound in RPT1, closing the nucleotide-binding pocket (Figure 5E and 5G). In contrast, the nucleotide-binding pocket of RPT5 gradually shrinks but is not fully closed by the R-fingers of RPT1, whereas the nucleotide-binding pocket of RPT4 gradually opens up (Figure 5G). As a result, the pore-1 loops of RPT1 and RPT5 are in contact with the substate in the last and first states E_D1.3_ and E_D0.1_, respectively. Both are dissociated from the substate in states E_D0.2_, E_D0.3_, E_D1.1_, and E_D1.2_. These gradual movements allow us to assign their temporal sequences as illustrated (Figures 5A–5E), indicating that these states represent intermediate conformations accompanying single-nucleotide exchange that occurs in RPT5 (Figure 5H). Interestingly, the outward flipping of RPT5 appears to complete its major conformational change in state E_D0.3_, much faster than the inward motion of RPT1, which completes its conformational change in state E_D1.3_ (Figure 5F). This observation suggests that ATPase re-engagement with the substrate is the rate-limiting step in the single-nucleotide exchange kinetics of the AAA-ATPase motor.

Our data clarify that the CP gate is open once the pore-1 loop of RPT2 binds the substrate in state E_D0.1_, as five RPT C-terminal tails are inserted in the α-pockets in all six states (Figure S3C). Altogether, these structures vividly characterize at the atomic level how the “hand-over-hand” actions of RPT subunits are executed by coordinated conformational dynamics in the AAA-ATPase motor during single-nucleotide exchange and generates mechanical force unfolding the substrate. Although the sequential model of coordinated ATP hydrolysis around the AAA- ATPase ring has been hypothesized in numerous similar systems,^2,6,7,12^ our results provide the first direct evidence for sequential intermediate states of hand-over-hand actions within a single cycle of nucleotide exchange.

### Hierarchical allosteric regulation of proteolytic activity

Both atomic structures of the substrate-bound CPs in the open- and closed-gate states E_A_ and E_D2_ show a segment of substrate polypeptide binding to the catalytically active residue Thr1 in the β2 subunit, whereas no substrate density is observed in the other two catalytic subunits β1 and β5. In both states, the substrate is bound in a β-sheet conformation between residues 20 to 22 and 46 to 48 in the β2 subunit (Figures 6A, 6B, S8A and S8B), reminiscent of the conformations of CP inhibitors binding to the β5 subunits in the crystal structures of the CP.^32,33^ The hydroxyl oxygen atom of the catalytic Thr1 is approximately 2-Å closer to the substrate mainchain in state E_D2_ compared to state E_A._ This structural difference is expected to result from the CP gate opening, which exerts a profound allosteric impact on the entire CP structure.^34–37^

**Figure 6.**
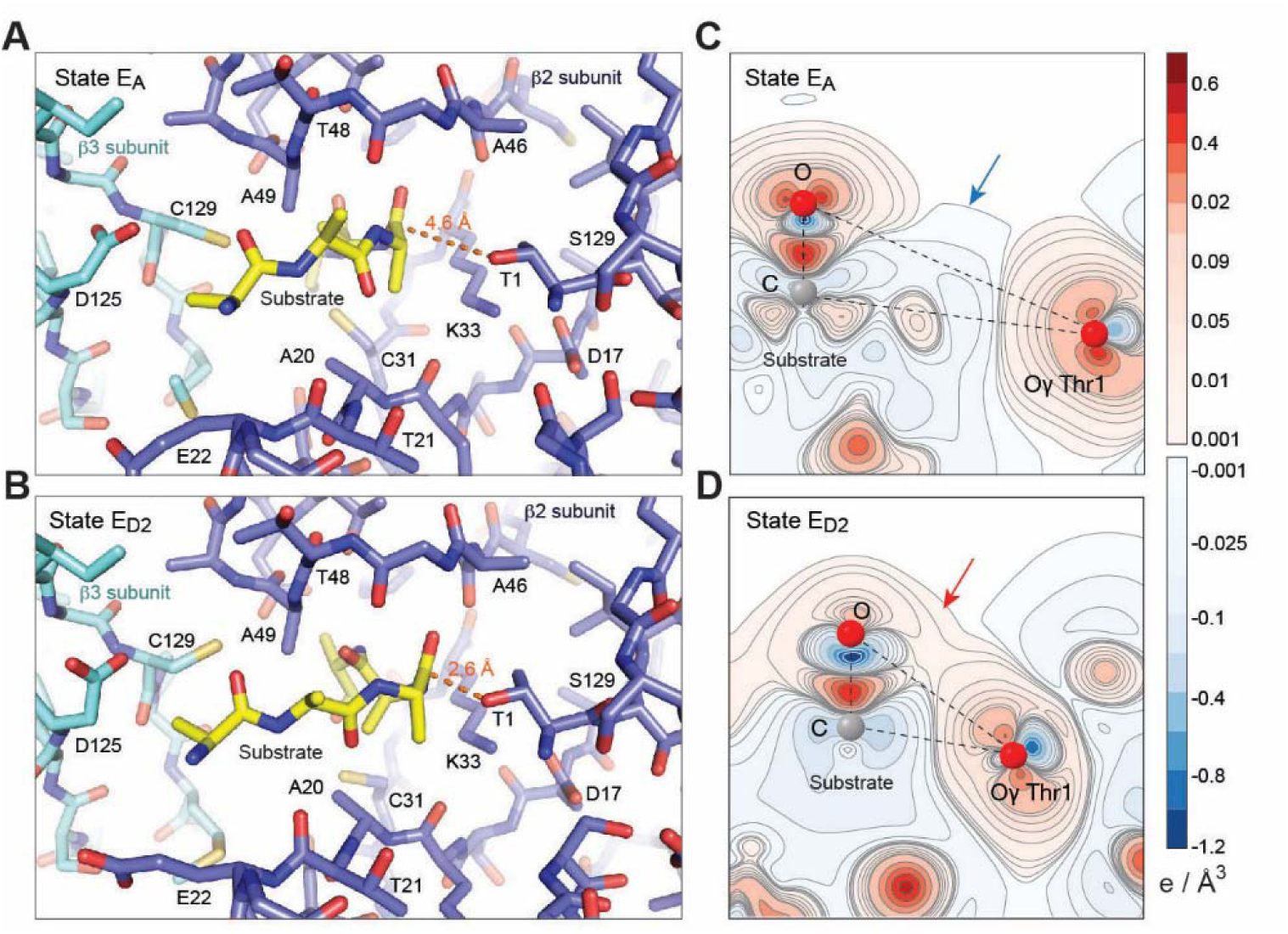
Allosteric regulation of substrate proteolysis by the opening of CP gate. (**A** and **B**), Structure of the catalytic site with a substrate polypeptide bound at residue Thr1 in the β2 subunit of the CP in state E_A_ at 2.7-Å resolution (**A**) and in state E_D2_ at 2.5-Å resolution (**B**). (**C** and **D**) Charge density difference contour maps at the proteolytic active site of the β2 subunit computed by the quantum mechanical DFT based on the atomic structures of states E_A_ (**C**) and E_D2_ (**D**). The contours are plotted in the plane passing through the Thr1-Oγ atom and the nearest carbonyl carbon and oxygen atoms of the substrate. Blue and red arrows highlight a non-bonding charge depletion region in state EA and a bonding interaction indicated by the slight positive charge difference region in state E_D2_, respectively.

To understand how such a subtle structural change influences proteolytic activity, we carried out first-principles calculations of the quantum-mechanical interactions involved in the nucleophilic attack of the catalytic residue Thr1 on the scissile peptide bond of the substrate, using the density functional theory (DFT) with the generalized gradient approximation.^38^ To initiate the nucleophilic attack, the carbonyl group must be brought into proximity with the electrophile Thr1-Oγ to facilitate formation of a tetrahedral transition state.^33,39,40^ We calculated the charge density difference induced around the interacting pair after proton transfer from Thr1-Oγ to Thr1-NH_2_ (Figures 6C, 6D, S8E and S8F). The substrate-interacting pair of atoms in state E_D2_ form a bonding region of slight positive charge difference linking the carbonyl group of the substrate to Thr1-Oγ. In contrast, the pair are separated by a charge depletion non-bonding region in state E_A_. Indeed, a recent study has found that the nucleophilic attack is the rate-limiting step in the acylation barrier.^39^ Our result suggests that CP gate opening allosterically promotes the nucleophilic attack by lowering the activation energy barrier of acylation and induces hybridization of molecular orbitals, hence upregulating the reaction rate of proteolysis. In support of our structural finding, previous studies have observed indirect evidence for coupling of the CP gating with the proteolytic activity.^34–37^

## Discussion

By analyzing 64 conformers under a common biochemical condition systematically, this study reveals a new level of complexity of the proteasome regulation. The previously underappreciated high degree of compositional and conformational heterogeneity of substrate-engaged 26S proteasome has not been similarly explored in other macromolecular machinery systems. The improved accuracy in 3D classification via our data-driven deep learning approach^17^ allowed us to discover many previously ‘invisible’ conformers and novel structural features, including additional ubiquitin-like densities, continuous conformational transitions of the AAA- ATPase motor driven by single-nucleotide exchange, and ATP-dependent allosteric regulation of substrate proteolysis in the β2 subunit. We noted several related recent methodological efforts in leveraging deep learning^41,42^ to analyze continuous protein motions from heterogeneous cryo-EM datasets, which nonetheless fell short of sufficient resolution in visualized conformers. In contrast to our approach, applications of these other related methods^41–43^ to the same dataset in this study did not recover the hidden conformers and failed to gain insights beyond previously reported results.^6^ Some difficulties originated from the way we prepared the cryo-EM sample using time-dependent ATP-to-ATPγS exchange, which was aimed at stalling the proteasome in a maximal number of intermediate conformations before the completion of substrate degradation.^6^ Most of the intermediate conformers were in thermodynamic states far away from equilibrium. This study underscores the importance of ultra-deep, highly accurate 3D classification of nonequilibrium conformers in understanding the functional dynamics of protein machinery as opposed to the traditional paradigm of cryo-EM analysis that is largely limited to solve a few equilibrium conformers. In a recent study, we systematically compared the intermediate conformers of substrate-engaged USP14-proteasome treated with ATP-to-ATPγS exchange to those with only ATP in the buffer.^8^ We observed only minor structural variations in the lid subunits in contact with the ATPase, such as the N-terminal regions of RPN5 and RPN6, but not in the structures of the ATPase motor.^8^ Thus, we do not expect that any of the newly reconstructed conformers to be non-physiological dead ends due to the use of ATP-to-ATPγS exchange protocol.

### Insights into ubiquitin-decoding architecture of the proteasome

By conducting complementary experimental examinations including mutagenesis, MST and LC-MS/MS, we confirm that RPN2 and α5 harbor low-affinity binding sites for ubiquitin. Our results provide insights into how the proteasome may interact with substrate-conjugated ubiquitin chains beyond the ‘canonical’ ubiquitin receptors. Both ubiquitin-binding sites on RPN2 and α5 are located in the vicinity of primary receptor sites. The RPN2 T1 site is very close to the RPN10 UIMs, whereas the α5 site is near the RPN1 T2 site. Since ubiquitin is also bound to RPN11 in states E_A2.2_, and E_B.2_, a Lys63-linked triubiquitin chain can be fitted into the RPN2- and RPN11-bound ubiquitin densities by modeling the rotation of Lys63 linkages. This suggests that the RPN2 T1 site may function as a coreceptor site for ubiquitin transfer from RPN10 to RPN11 and facilitate deubiquitylation (Figure 7A). Given that at least one RPN10 UIM-bound ubiquitin invisible in these cryo-EM maps is expected to exist, the observation of triubiquitin on RPN2 and RPN11 structurally rationalizes why tetraubiquitin is an optimal degradation signal for the proteasome^2^. Interestingly, atomic structure modelling of a Lys63-linked tetraubiquitin chain by fitting its terminal ubiquitin into the cryo-EM density suggests that a single tetraubiquitin can span across the α5 subunit and the RPN1 T2 site (Figure 7B). Since RPN1 appears to allosterically regulate the AAA-ATPase motor conformation^6^ (Figures 5A and 5B), tetraubiquitin binding to both the RPN1 T2 site and α5 subunit could help to allosterically stabilize the open-CP states (E_D0_, E_D1_ and E_D2_).

**Figure 7.**
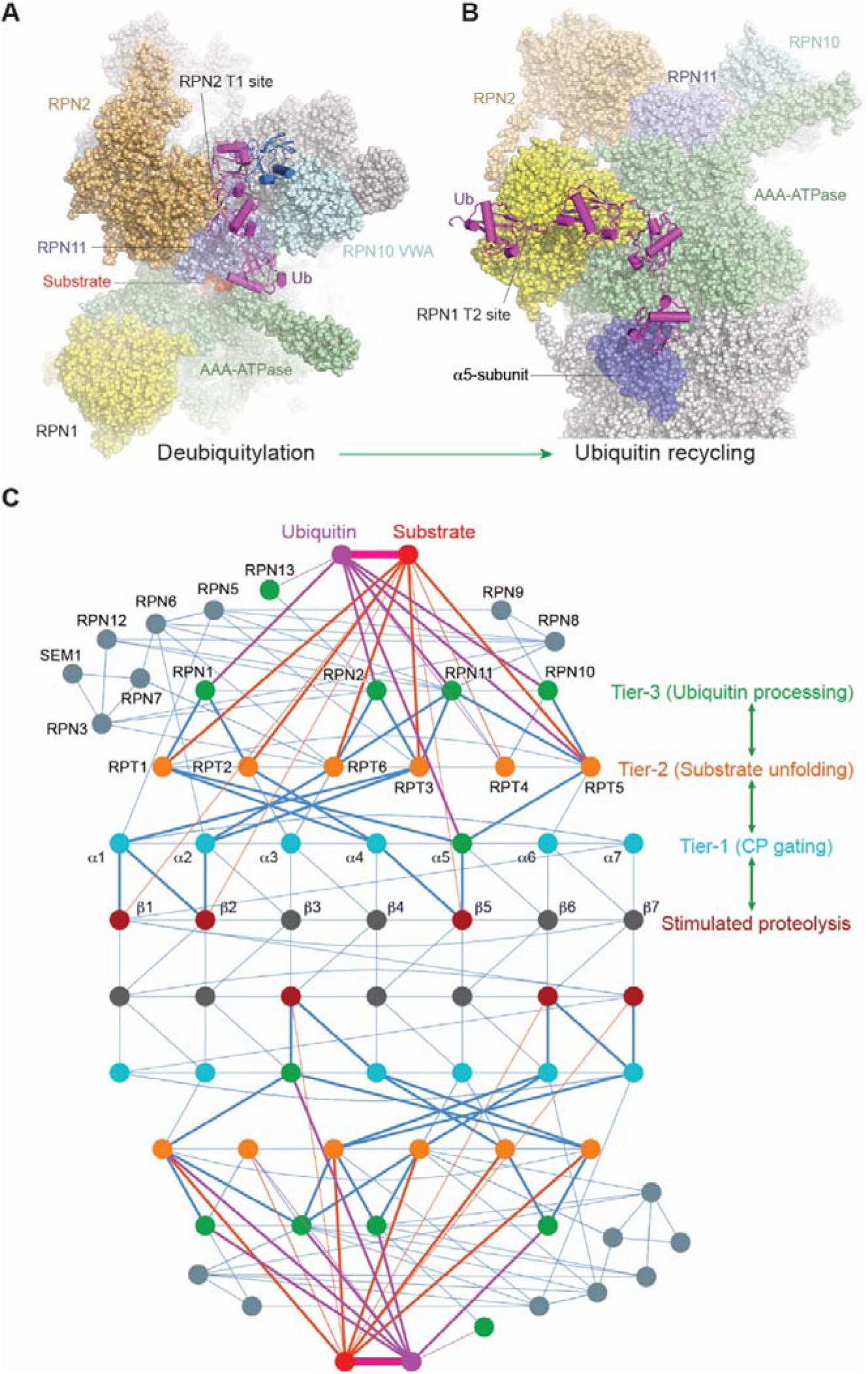
Insights into ubiquitin-mediated autoregulation of the human 26S proteasome. (**A**) A Lys63-linked triubiquitin chain modelled from the cryo-EM maps of states E_A2.2_ and E_B.2_ is shown in purple cartoon representation and spans from the RPN2 T1 site to the RPN11 deubiquitylation site. The RPN10 VWA-bound monoubiquitin fitted in states E_A2.3_ and E_B.3_ is superimposed in blue cartoon representation. The proteasome is shown as space-filling sphere representations. (**B**) A Lys63-linked tetraubiquitin chain model derived from the cryo-EM map of state E_D2.2_. The two terminal ubiquitin molecules are fitted from their highest-resolution density maps available at the RPN1 T2 and α5-subunit sites, whereas the middle two molecules are hypothetically modelled. (**C**) A network model of hierarchical allosteric regulation of proteasome activity. The interactome network diagram illustrating the grand allostery of the proteasome. Steel blue lines represent inter-subunit interactions; purple lines represent the interactions with ubiquitin; and red lines represent substrate polypeptide interactions. Bold lines highlight the shortest pathways connecting the three tiers of regulatory subunits that are expected to propagate the allosteric regulation from ubiquitin interactions down to the six proteolytic active sites in the CP. The network diagram is constructed according to all 64 structural models of the substrate-bound proteasome in this study. Notably, the subnetwork architecture of RPN1 connecting to the proteolytic subunits is identical to that of RPN2 connecting to the proteolytic subunits, consistent with the structural homology of RPN1 and RPN2 as well as their potential functional similarity in ubiquitin interactions.

The ubiquitin code targeting substrates to the proteasome primarily consist of Lys48-linked polyubiquitin chains.^44^ Proteomic and biochemical analyses confirm that other types of linkages are also used for substrate targeting by the proteasome, including Lys63 linkage that were initially thought to mediate non-proteasomal pathways.^45,46^ Certain linkage types including hybrid Lys48/Lys63-linked branched chains may be more or less efficient in mediating proteasomal degradation by enhancing ubiquitin-chain recognition or triggering efficient, rapid ubiquitin removal via deubiquitylating enzymes (DUBs) such as USP14/Ubp6 and UCHL5/Uch37. Deubiquitylation prior to substrate commitment to the proteasome prevents substrates from being degraded by the proteasome, thus stabilizing their intracellular protein level. The low-affinity ubiquitin-binding sites on RPN2 or the α5 subunit alone are expected to be insufficient in primary targeting of substrates to the proteasome.

However, it is conceivable that, after the high-affinity sites first capture certain polyubiquitin chains, binding of adjacent polyubiquitin chains to the nearby low-affinity sites could facilitate recognition of hybrid or branched ubiquitin chains. Stepwise binding of ubiquitin moieties might be kinetically important for the proteasome in recognizing branched ubiquitin chains due to kinetic heterogeneity in recognizing difference ubiquitin chains. This mechanism could underline the phenomenon that certain branched ubiquitin chains can promote proteasomal targeting of specific substrate proteins.^45,46^ The ubiquitin-binding sites of both high and low affinity together could remodel multiple polyubiquitin chains simultaneously, deliver the peptide-proximal ubiquitin to the deubiquitylation site in time (Figure 7A), as well as assist in ubiquitin recycling after hydrolysis of isopeptide bonds (Figure 7B).^6,8^

### A refined mechanism for substrate translocation

The six conformers visualizing single-nucleotide exchange dynamics allows us to elaborate on an improved mechanistic understanding of substrate unfolding and translocation. Recent years have seen cryo-EM analysis of numerous substrate-engaged ATPase translocase complexes from AAA+ superfamily^47^. Proteasomal ATPases belong to the classic clade of AAA+ superfamily and thus could potentially employ a mechanism of substrate translocation that is also assumed in other related clades. Indeed, most currently available cryo-EM structures of substrate-engaged AAA+ ATPase complex share a highly similar architecture of substrate-pore loop interactions as those observed in the proteasome.^6–8^ These studies consistently proposed a hypothetical hand-over-hand model of substrate translocation, despite lack of detailed evidence for how the hydrolysis and exchange of nucleotide drive the motorization of the ATPase complex within a cycle of single-nucleotide exchange.^47^

The initial proposal of hand-over-hand model implies that sequential hydrolysis of ATP around the ATPase ring is structurally and dynamically symmetric in typical AAA+ ATPase homohexamers like Yme1 and ClpXP.^47^ By contrast, the asymmetric architecture of substrate-engaged ATPase heterohexamer in the proteasome underlies its functional asymmetry.^8,48–50^ Growing evidence points to the asymmetric interactions of the ATPase motor with various proteasome-associated factors— including USP14/Ubp6 and ZFAND5—that regulate differential activation of the proteasome^8,51,52^. Notably, the six conformers of the substrate-engaged ATPase ring within a cycle of single-nucleotide exchange resolved in this work indicate two consecutive kinetic sub-steps: (1) faster dissociation of the ADP-releasing ATPase subunit from the forefront of substrate movement near the CP gate and (2) slower re-association of the ATP-exchanged ATPase subunit with substrate near the entrance of the OB ring. Consistently, recent computational modelling of the proteasome found such kinetic sub-steps to be crucial in recapitulating the dynamics of substrate translocation *in silico*.^53^ Our observation of differential substrate movement toward the CP gate excludes the possibility of a single-step, burst translocation model recently hypothesized for the ClpXP degradation complex, which operates in the opposite direction of ATP hydrolysis around the ring.^54^

Taking into account all existing evidence from this and related studies,^6,8,53^ our reconstitution of single-nucleotide exchange dynamics of the proteasomal ATPase motor refines the mechanism of substrate translocation. We postulate that the kinetic variation of the second sub-step, namely, the step of slow re-association of the ATP-exchanged ATPase subunit with the substrate, can accommodate certain probabilistic events such as substrate sliding or slipping. During this transient interval, two or more substrate-dissociated ATPase subunits would result in a weaker gripping of substrate that permits substrate translocation driven by its own conformational entropy. This model revises the strictly sequential hand-over-hand mechanism by incorporating a sub-step of probabilistic sliding of substrate during the interval when the fewest ATPase subunits grip the substrate. This refined model can potentially reconcile the cryo-EM observation of two residues gripped between adjacent pore loops and other experimental observation of substrate translocation of more than two residues per ATP hydrolyzed.^54^ We anticipate that further studies testing this refined model will help establish a universal mechanistic framework for AAA+ family translocases.

### An integrative allostery model for proteasome autoregulation

Our systematic structural analyses posit a grand allostery model for proteasomal autoregulation of proteolytic activity operating through a three-tier hierarchy of inter-molecular interaction networks (Figure 7C), where all subunit conformations are allosterically coupled. The first tier of autoregulation exists between the CP gate in the α-ring and the catalytic sites of the β-subunits. Opening of the CP gate allosterically upregulates the proteolytic activity of the three β-subunits. The second tier of autoregulation lies between the substrate-bound AAA-ATPase motor and the rest of the proteasome. The AAA-ATPase motor bridges the ubiquitin-binding sites and the CP via direct interactions of the RPT coiled-coil (CC) domains with RPN1, RPN2 and RPN10, which harbor ubiquitin-binding sites. As a result, three modes of coordinated ATP hydrolysis appear to regulate intermediate steps of substrate processing^2,6^, with initiation of substrate unfolding allosterically coupled to CP gating during E_C2_ to E_D0.1_ transition (Figure 5). The third tier of autoregulation involves dynamic interactions between ubiquitin signals and a collection of ubiquitin receptor sites and DUBs. This is manifested in the dual roles of ubiquitin-binding sites in recognizing ubiquitin signals and in transmitting such recognitions to allosteric regulation of AAA-ATPase conformations that controls the CP gate. Ubiquitin binding and remodeling destabilize the resting state of the proteasome, promoting its transition to states E_B_, E_C_ and ultimately the E_D_-like states. On the other hand, a polyubiquitin chain binding multiple sites within the proteasome can exert profound allosteric effects on conformational selection (Figures 1G and 1H). Such three-tier, hierarchical allosteric regulations may occur mutually with both RPs and are expected to underlie the conformational entanglement of the two RPs in the DC proteasome (Figure 1E), which was not clearly observed in the absence of substrates.^2,13,14^ This grand allostery mechanism might have been evolved to optimize proteasome function in response to extremely complex intracellular challenges and is manifested in various phenotypic effects, such as UBL-induced proteasome activation^55,56^ and long-range allosteric couplings in the proteasome.^2,35,36,57,58^

### Limitations of the study

The current study used substrate mainly modified by Lys63-linked ubiquitin chains. This limited the structural characterization of ubiquitin chain interactions with the proteasome. Replacing the model substrate with Lys48-linked chains or with other linkage types including branched chains is anticipated in future studies. Due to the limited resolution of ubiquitin and transient nature of these ubiquitin-bound states, we cannot entirely rule out the possibility that some of the ubiquitin-like densities correspond to co-purified endogenous regulators of the proteasome. The absence of other cellular regulators of the proteasome in our reconstituted degradation system, such as UBE3A, UBE3C, ZFAND5 and ECM29, limited our understanding of the proteasome regulation under other various physiological conditions. Future investigations are needed to address these challenges and to reveal the underexplored aspects of proteasome function and regulation.

## STAR Methods

### Protein purification and sample preparation

Human 26S proteasomes were affinity-purified as previously described.^6^ In brevity, a stable HEK293 cell line expressing HTBH (hexahistidine, TEV cleavage site, biotin, and hexahistidine)-tagged hRPN11 was used to purify the human proteasome on a large scale.^25^ After Dounce-homogenization in a lysis buffer (50 mM PBS [pH 7.5], 10% glycerol, 5 mM MgCl_2_, 0.5% NP-40, 5 mM ATP, 1 mM DTT, and protease inhibitors), cell lysates were cleared by centrifugation (20,000 × g) for 30 min. The supernatant was incubated with NeutrAvidin agarose resin (Thermo Scientific) for 3 h at 4 °C. 26S proteasomes were cleaved from the agarose beads by TEV protease (Invitrogen) after washing with excess lysis buffer and a wash buffer (50 mM Tris-HCl [pH 7.5], 1 mM MgCl_2_ and 1 mM ATP). The supernatant was then further purified by gel filtration on a Superose 6 10/300 GL column at a flow rate of 0.15 ml/minute in buffer (30 mM Hepes [pH 7.5], 60 mM NaCl, 1 mM MgCl_2_, 10% glycerol, 0.5 mM DTT, 0.6 mM ATP). Gel-filtration fractions were concentrated to about 2 mg/ml and the buffer was exchanged to 50 mM Tris-HCl (pH 7.5), 100 mM NaCl, 1 mM ATP and 10% glycerol.

The model substrate Sic1^PY^ was expressed in BL21 (DE3) *Escherichia coli* cells and purified as previously described.^6^ The PY motif, Pro-Pro-Pro-Tyr, is recognized by WW domains the Rsp5 family of ligases. In the Sic1^PY^ construct, a PY motif was inserted to the N-terminal segment (MTPSTPPSRGTRYLA) of the Cdk inhibitor Sic1, resulting in a modified N-terminus of MTPSTPP<u>PPY</u>SRGTRYLA.^59,60^ The ubiquitylation reaction of Sic1^PY^ was conducted by mixing 40 μg/ml Sic1^PY^ with 500 nM E1 Ube1 (Boston Biochem), 2 μM E2 UbcH5a (Boston Biochem) and 100 μg/ml E3 WWHECT and 1 mg/ml ubiquitin (Boston Biochem) in a reaction buffer (50 mM Tris-HCl [pH 7.5],100 mM NaCl, and 10% glycerol, 2 mM ATP, 10 mM MgCl_2_, and 1 mM DTT) for 3 h at room temperature. The recombinant E3 ligase WWHECT was engineered from deletion of Rsp5 N-terminal 220 amino acids, which conjugates ubiquitin chains via Lys63 linkage.^61^ Pre-equilibrated TALON resin was then incubated with the sample for 1 h at 4°C. After the resin was washed with 20 column volumes of the wash buffer (50 mM Tris-HCl [pH 7.5], 100 mM NaCl, 10% glycerol), the polyubiquitylated Sic1^PY^ was eluted with the same buffer containing 150 mM imidazole. The elution was applied to an Amicon ultrafiltration device with 30K molecular cut-off for removal of imidazole. The ubiquitylation results were examined by Western blotting with anti-T7 antibody.

### Liquid chromatography–tandem mass spectrometry

Liquid chromatography–tandem mass spectrometry (LC–MS/MS) assay was used to analyze the ubiquitination of endogenously purified 26S proteasome subunits. Briefly, 26S proteasomes were incubated with 5 mM DTT and 10 mM iodoacetamide (IAA) for 1 h, respectively, and then digested with trypsin overnight. The enzymatic hydrolysis was terminated by 0.1% trifluoroacetic acid (TFA) and subjected to MS analyses. LC-MS/MS analysis were performed on an Easy-nLC 1200 system connected to Thermo Orbitrap Fusion Lumos (Thermo Fisher Scientific). Peptide mixtures were loaded onto a C18 column (Acclaim PepMap 100, 75 μm × 2 cm nanoViper Thermo) at a speed of 280 nL/min and separated using a segmented gradient in 75 min. Solvent A consisted of 0.1% (v/v) formic acid in H_2_O and solvent B consisted of 0.1% (v/v) formic acid in 80% acetonitrile. The gradient was set as follows: 4%–8% solvent B in 5 min; 8%–20% solvent B in 45 min; 20%–30% solvent B in 10 min; 30%–90% solvent B in 13 min; 90% solvent B in 3 min. The mass spectrometer was operated in a data-dependent mode. Full MS scans were conducted using an Orbitrap Mass Analyzer at the range of 300-1500 m/z with the resolution set to 120K at *m/z* 200, automatic gain control (AGC) target at 5 × 10^5^, and a maximum injection time at 90 ms, followed by MS/MS scans generated by HCD (30% energy) fragmentation at the resolution of 30K, AGC target 5 × 10^4^, and a maximum injection time of 100 ms. All data generated were searched against the Uniprot *Homo sapiens* (Human) using Proteome Discoverer v.2.2 software (Thermo Fisher Scientific). Search criteria used were trypsin digestion, allowing up to 2 missed cleavages, mass accuracy of 10 ppm for the precursor mass tolerance and 0.02 Da for the fragment ions mass tolerance. Carbamidomethylation (C) was set as fixed modification and Ubiquitination (K), Oxidation (M) were set as variable modification. The false discovery rate (FDR) applied at the protein-level and peptide-level was 1%.

### Site-directed mutagenesis of RPN2 and **α**5

Site-directed mutagenesis of RPN2 (Q527A/G534R/E538R in the “RPN2 T1 site”) is based on the site-directed mutagenesis of RPN1 T1 site and the atomic model of state E_D2.2_, as structural and electrostatic properties of RPN1 and RPN2 T1 sites are highly conserved (Figures 2D, S3H–S3K). The sequence of mutant RPN2 genes and primers was synthesized by Genscript (Nanjing, China) and cloned into pGEX-6P vector. Site-directed mutagenesis of α5 is based on the atomic model of state E_D2.2_. According to the atomic model of state E_D_2.2, the ubiquitin is near to the loop connecting two helices (Ser174-Ser180 and Leu191-Gln204) on α5 subunit, which contains acidic or basic residues Glu183, His186 and Lys187. These acidic or basic residues may have ionic interaction with residues on ubiquitin. Therefore, we performed triple site-directed mutagenesis (E183A/H186A/K187A) on the α5 subunit to verify the ubiquitin-binding site. The sequence of mutant α5 genes and primers was synthesized by Genscript (Nanjing, China) and cloned into pGEX-4T vector.

### Expression and purification of human RPN2 and its mutants

Genes for wild-type RPN2 and its mutants (Q527A, G534R, E538R) were cloned into pGEX-6P vector (GenScript, Nanjing), which contains a GST-tag and a HRV3C protease cleavage site at N-terminus, to get two plasmids. BL21-CondonPlus (DE3)-RIPL cells (Shanghai Weidi) transformed with the above two plasmids were grown to an OD_600_ of 0.6–0.8 in LB medium supplemented with 100 mg ml^−1^ ampicillin and then gently cooled to 16 °C. Cells were induced with 75 μM IPTG at 16 °C for 20 h and collected by centrifugation at 3,000g for 15 min. Cells were lysed in lysis buffer (50 mM Tris-HCl [pH 7.5], 150 mM NaCl, 1 mM DTT, 0.2% NP-40 and 1× protease inhibitor cocktail) and the lysate was cleared through centrifugation at 20,000g for 60 min at 4 °C. The supernatant was incubated with GSTSep Glutathione Agarose Resin (Yeasen) for 3 h at 4 °C. The resin was washed with 20 column volumes of washing buffer (50 mM Tris-HCl [pH 7.5], 300 mM NaCl and 1 mM DTT) and 20 column volumes of cleavage buffer (50 mM Tris-HCl [pH 7.5], 150 mM NaCl, 1 mM DTT and 10% glycerol). The GST tag was cleaved by incubating with HRV 3C Protease (Thermo Scientific) overnight at 4 °C in 1.5 ml cleavage buffer. The cleaved samples were further purified with anionic exchange column (HiTrap Q HP, cytiva) and gel filtration column (Superdex 200 10/300 GL, GE Healthcare) equilibrated with 50 mM Tris-HCl (pH 7.5), 100 mM NaCl, 1 mM DTT and 10% glycerol.

### Expression and purification of human **α**5 and its mutants

Wild-type α5 and its mutants (E183A, H186A, K187A) genes were cloned into pGEX-4T vector obtained from GenScript (Nanjing, China) to get two plasmids. To purify recombinant human Alpha5 and its mutant, BL21-CondonPlus (DE3)-RIPL cells (Shanghai Weidi) transformed with the above two plasmids were grown to an OD_600_ of 0.6 and then induced by 0.1 mM IPTG for 20 h at 16 °C. Cells were collected by centrifugation at 3,000× *g* for 15 min, resuspended in lysis buffer (50 mM Tris-HCl (pH 7.5), 150 mM NaCl, 0.2% NP-40, 1 mM DTT and 1× protease inhibitor cocktail) and lysed by sonication. The lysate was cleared through centrifugation at 20,000× *g* for 60 min at 4 °C. The supernatant was incubated with GSTSep Glutathione Agarose Resin (Yeasen) for 3 h at 4 °C. The resin was washed with 20 column volumes of washing buffer (50 mM Tris-HCl (pH 7.5), 300 mM NaCl and 1 mM DTT) and 20 column volumes of cleavage buffer (20 mM Tris-HCl (pH 8.0), 150 mM NaCl, 1 mM DTT). To cleave the GST tag, resin is incubated with thrombin (Solarbio) in 1.5 ml cleavage buffer for 2 h at 30 °C followed by overnight incubation at 4 °C. The cleaved samples were further purified with anionic exchange column (HiTrap Q HP, cytiva) and gel filtration column (Superdex 75 10/300 GL, GE Healthcare) equilibrated with 50 mM Tris-HCl (pH 7.5), 100 mM NaCl, 1 mM DTT and 10% glycerol.

### Microscale thermophoresis

The human RPN2/α5 and their mutants were labelled with the 2nd Generation dye of RED-NHS using the Monolith Series Protein Labeling Kit (NanoTemper). Before labelling, proteins were exchanged to the labeling buffer (provided in the kit) containing 130 mM NaHCO_3_ and 50 mM NaCl with column A (provided in the kit). After labelling, excess dye was removed by applying the sample on column B (provided in the kit) equilibrated with reaction buffer (50 mM Tris-HCl (pH 7.5), 100 mM NaCl, 1 mM DTT, 1 mM EDTA and 0.05% Tween-20). 0.05% Tween-20 was added to the sample before MST measurements. For interaction of RED-NHS-labelled RPN2/α5 (or their labelled mutants) with mono-ubiquitin, concentration series of mono-ubiquitin were prepared using a 1:1 serial dilution of protein in reaction buffer containing 0.05% Tween-20. The range of mono-ubiquitin concentration used began at 6 mM, with 16 serial dilution in 10-μl aliquots. The interaction was initiated by the addition of 10 μl of 1 μM RED-NHS-labelled RPN2/α5 (or their labelled mutants) to each reaction mixture and measured by Monolith NT.115 (NanoTemper) at 2% LED excitation power and 40% MST power.

### GST pull-down experiments

600 pmol of purified GST-tagged RPN2 full-length or 600 pmol of purified GST (negative control) was added to 50 μL of prewashed GSTSep Glutathione Agarose Resin (Yeasen) in 500 μL incubation buffer (50 mM HEPES (pH 7.0), 50 mM NaCl, 5% (v/v) glycerol, 0.2 mg/ml BSA, 1 mM EDTA and 2 mM DTT) for three hours and washed twice (1ml each time) with washing buffer (50mM HEPES (pH 7.0), 100 mM NaCl, 5% (v/v) glycerol, 0.2 mg/ml BSA, 1 mM EDTA and 2 mM DTT). The resin was next incubated with 10 pmol of M1-, K6-, K11-, K27-, K29-, K33-, K48-, and K63-linked diubiquitin in 100 μL incubation buffer above for one hour and washed twice (1ml each time) with same incubation buffer. Before elution, we washed the resin for pre-equilibrium with 1 ml pre-equilibrium buffer (50 mM HEPES (pH 8.0) 50 mM NaCl, 5% (v/v) glycerol, 1 mM EDTA and 2 mM DTT). Finally, the proteins retained on the pre-equilibrated resin were eluted by 60 μL elution buffer (30 mM glutathione, 50 mM Tris-HCl, 100 mM NaCl and 1 mM DTT at pH 8.0), fractionated by electrophoresis, transferred to a PVDF membrane and analyzed by immunoblotting.

### Homology sequence alignment of RPN2 and **α**5

Multiple sequence alignment (MSA) of RPN2 was performed for five species (six sequences, as *Arabidopsis thaliana* has two sequences: RPN2a and RPN2b). The alignment range for *homo sapiens* is 491-575, which was determined based on the RPN1 T1 site^11^ and the atomic model of state E_D2.2_. The alignment ranges for other species were determined by the model fitting used in Chimera (using command **mm** to fit two atomic models).

Similarly, MSA of α5 was performed across five species (six sequences, as *Arabidopsis thaliana* has two sequences: alpha type-5-A and alpha type-5-B). In the atomic model of state E_D2.2_, the ubiquitin is in proximity to the loop (181-190) connecting two helices (Ser174-Ser180 and Leu191-Gln204) on the α5 subunit. Therefore, the alignment range for Homo sapiens was selected as 181-190 (Figure 3C). The alignment ranges for other species were also determined by the model fitting used in Chimera.

### Cryo-EM data collection

The purified proteasome (10 μl of 2 mg/ml) was mixed with polyubiquitylated Sic1^PY^ (9 μl of 2 mg/ml; molar ratio ∼3:1 for substrate vs proteasome) in a buffer (50 mM Tris-HCl [pH 7.5], 100 mM NaCl, 10% Glycerol and 1 mM ATP) at room temperature. After 30 s of incubation of the mixture, 1 μl 20 mM ATPγS was immediately added into the solution to quench the degradation reaction. To remove the glycerol, the complex system was applied to Zeba™ Micro Spin Desalting Columns (7K, Thermo Fisher Scientific, MA, USA), exchanging the buffer to 50 mM Tris-HCl (pH 7.5) containing 100 mM NaCl, 1mM ATP and 1 mM ATPγS. The glycerol removal process usually took about 10 min before cryo-plunging. 0.005% NP-40 was added to the proteasome solution immediately prior to cryo-plunging. A 2.5-μl drop of 1.5 mg/ml substrate-engaged proteasome solution was applied to the glow-discharged C-flat grids (R1/1 and R1.2/1.3, 400 Mesh, Protochips, CA, USA) in a Vitrobot Mark IV (Thermo Fisher Scientific, MA, USA) chamber with 100% humidity at 4 °C. After 2 s of blotting, the grid was plunged into liquid ethane and then transferred to liquid nitrogen. Cryo-EM images were collected on an FEI Titan Krios G2 microscope equipped with the post-column Gatan BioQuantum energy filter connected to Gatan K2 Summit direct electron detector. Coma-free alignment was manually optimized and parallel illumination was examined prior to data collection. The data collection was semi-automatically executed by Leginon^62^ version 3.1 and SerialEM^63^ with the Gatan K2 Summit operating (Gatan Inc., CA, USA) in a super-resolution electron-counting mode and with the Gatan BioQuantum operating in the zero-loss imaging mode. Each movie contains 40 frames recorded with a total exposure time of 10 s with 250 ms per frame, with an accumulated dose of 44 electrons/Å^2^. The calibrated physical pixel size and the super-resolution pixel size are 1.37 Å and 0.685 Å, respectively. The defocus was set in the range of -0.7 to -3.0 μm during data collection of a total of 44,664 movies. Cryo-EM data pre-processing, including motion correction, micrograph screening, and particle picking by deep-learning-based algorithm, was conducted as previously described.^6^

### Analysis of substrate-engaged proteasome dataset

Prior to 3D classification by the procedure of AlphaCryo4D, only 2D classification by ROME^64^ and removal of apparent contaminants were conducted as previously reported^6^. All particles were binned twice down to 300 × 300 pixels prior to data processing. The general procedure of 3D classification by AlphaCryo4D has been described in detail elsewhere^65^, which includes the steps of *M*-fold particle shuffling and Bayesian resampling of volume data, deep manifold learning of energy landscape, and energy-based particle voting. To proceed with AlphaCryo4D, the whole dataset was randomly split into 32 sub-datasets, each with 100,000 particles. 3-fold particle shuffling and volume resampling procedures were conducted on each sub-dataset, which enhanced the total data size to 3,254,352 × 3 particles and yielded 32 × 4 × 10 = 1280 volumes in total that were used to compute the energy landscape of the substrate-engaged proteasome (Figures 1B and 1C). Using all available particles after manifold learning in AlphaCryo4D, the free energy value of each point in the landscape was calculated using the following equation:

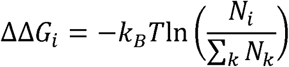

where ΔΔG_*i*_ denotes the free energy difference of a data point with the particle number of N_*i*_ against a common reference energy level, k_*B*_ is the Boltzmann constant and T is the temperature in Kelvin. By substituting the corresponding particle numbers, we generated landscapes in the same t-SNE coordinate system as shown in Figure 1A-C. Particle-voting on this energy landscape detected previously missing states (such as E_D0.2,_ E_D0.3_, E_D1.1_ and E_D1.2_, etc.) and yielded previously determined states with improved map quality, such as E_A2_ and E_D2_. Guided by this initial discovery of new conformational states, extensive focused 3D classifications in AlphaCryo4D were performed, as described in detail in the next section, to visualize novel ubiquitin-binding sites and hidden dynamics of proteasomal AAA-ATPase motor during single-nucleotide exchange. The Bayesian clustering during the resampling step and the final refinement of classified particles were conducted in RELION.^15^

### Analysis of substrate-free proteasome dataset

The dataset of the ATPγS-bound substrate-free human 26S proteasome^14^ contains 455,680 particles and was previously used to determine six coexisting conformational states of the substrate-free proteasome by AlphaCryo4D using the same procedure as that for the dataset of the substrate-engaged proteasome.^17^ The cryo-EM data were recorded with the super-resolution pixel size of 0.75 Å. The dataset is publicly available in the Electron Microscopy Public Image Archive (EMPIAR) database at EMDataResource.org (accession code EMPIAR-10090). This dataset was subject to re-analysis by AlphaCryo4D procedure. The size of ATPγS-bound proteasome dataset was enhanced to 455,680 × 3 particles and was divided into 4 sub-datasets for volume resampling. 160 volumes were resampled to compute the energy landscape the substrate-free ATPγS-bound proteasome (Figure 1A), following the same protocol of AlphaCryo4D^17^ as that used for the analysis of the substrate-engaged proteasome dataset described in this paper.

### Focused 3D classification and reconstruction

To obtain a more complete picture of the conformational states of the substrate-engaged proteasome, we conducted focused 3D classification using AlphaCryo4D on several local regions on the overall energy landscape computed in the previous steps of analysis. To study the local features of ubiquitin binding and substrate interactions, 3D masks corresponding to the local densities of interest were applied to all resampled volume data, and the subset of interest on the energy landscape was extracted to conduct a zoomed-in analysis. In this case, the energy landscape calculated via masked volumes is expected to reflect the local structural variations of interest (Figure S2).

To detect novel ubiquitin-binding sites in the proteasome, local masks were applied to focus on the features around the putative locations of ubiquitin chains. After particle shuffling for *M* = 3 folds, 732,666 × 3 particle images of the state E_A_ and E_B_ were utilized for computing the zoomed-in energy landscape with 280 volumes. These particles were first aligned in a consensus alignment without any mask or resolution limit to restrict alignment parameter calculations. On the focused energy landscape, only the region within the soft mask of the ubiquitin and its binding site was applied in the volume resampling step, with the signals of the CP of the proteasome subtracted from each raw particle when dealing with the state E_A_ and E_B_. Accordingly, the particle diameter applied in the CP subtracted proteasome was shrunk to 274 Å, instead of 411 Å in the complete 26S proteasome. When the region around RPN2 masked for focused 3D classification to search for potential ubiquitin-like features, the total particle numbers were enhanced to 208,700 × 3 (E_A1_), 240,475 × 3 (E_A2_), 260,021 × 3 (E_B_) and 129,492 × 3 (E_D2_). The energy landscapes were computed with 80, 80, 80 and 40 masked volumes resampled by upper resolution bounds of 15, 15, 15 and 20 Å, respectively. Several 3D classes with the particle numbers of 192,219, 2,539, 3,982 and 609 were obtained by the particle- voting procedure on the zoomed-in energy landscape of state E_A1_, while the case of state E_A2_ gave four classes containing 147,108, 5,842, 5,500 and 33,083 particles (Figures S2F and S2G). Likewise, the particle numbers of 3D classes on the zoomed-in energy landscape of state E_B_ were 173,931, 5,117 and 9,754 (Figure S2H); and the zoomed-in energy landscape of state E_D2_ resulted in four classes having 61,580, 6,192, 24,674 and 1,673 particles, respectively (Figure S2I).

To detect and improve RPN1-bound ubiquitin densities, a local mask around RPN1 was used, with 15 Å resolution limit applied in the volume-resampling step. The zoomed-in energy landscape around state E_A1_ computed from 120 volumes resulted in five classes with the particle numbers of 128,161, 7,754, 6,557, 6,826 and 3,937. Five 3D classes on the zoomed-in energy landscape of E_A2_, also computed with 120 volumes, including 81,612, 27,483, 12,247, 26,029 and 12,777 particles, were analyzed (Figures S2C and S2D). Based on the subclass of E_A1_ that contains 128,161 particles, we continued to refine the alignment locally to the component of RPN1. Then a 3D mask around RPN1 was applied to resample 40 volumes with 20 Å resolution limit in the expectation step. After voting on the energy landscape, two classes with the particles number of 43,957 and 43,620 were generated, one of which exhibited a ubiquitin-like density on the T1 site of RPN1 and was designated state E_A1.1_ (Figure S2E). In the case of the ubiquitin-binding site on the α5 subunit of the CP, five 3D classes with the particle numbers of 136,071, 19,351, 4,100, 131,570 and 4,932 were generated by particle voting on the zoomed-in energy landscape of E_D_ computed with 120 volumes, which were resampled from 326,409 × 3 particles via the mask containing this ubiquitin-binding site and no high-resolution restriction (Figure S2J). During these analyses, all masked volume-resampling steps were performed for 50 iterations, whereas unmasked 3D classifications were performed for 30 iterations with the initial reference low-pass filtered to 60 Å. At the last step, 3D refinement and reconstruction of each class was done to finalize the high-resolution structure determination.

To exploit intermediate conformations during the substrate translocation initiation, a zoomed-in energy landscape was computed for the focused 3D classification procedure. First, the energy landscape of all 3,254,352 × 3 particles was computed and analyzed. Then the zoomed-in energy landscape of the states E_C_ and E_D_ containing 2,521,686 × 3 particle images were computed from 1,000 volumes that were resampled without masking or resolution restriction (Figure S2B). For further zoomed-in processes, the particles around the translocation initiation states (from E_C_ to E_D0_) on this energy landscape were extracted by the distance-based 3D classification strategy. Subsequently, these particles were shuffled 3 times to the number of 410,098 × 3 to generate 160 volumes for the computation of the zoomed-in energy landscape with the RP mask and no resolution restriction imposed in the volume resampling, which led to the 3D classes E_C1_, E_C2_, and E_D0.1_ with the particle number of 68,506, 77,545, and 25,755, respectively (Figure S2K). Moreover, state E_D0.3_ came from two clusters with 86,415 and 83,107 particles merged together on the unmasked energy landscape, and the classes E_D0.2_ and E_D1.1_ including 105,081 and 51,808 particles, respectively, were also obtained after particle voting on this energy landscape. In the local areas of the state E_D1_ on the energy landscape, two clusters with 80,841 and 94,273 particles were merged to reconstruct the high-resolution structure of state E_D1.2_, while states E_D1.1_ and E_D1.3_ with 40,696 and 145,506 particles, respectively, were separated out from the clusters on this unmasked energy landscape. In these studies, the maximum number of iterations in the 3D classifications of volume resampling was 30, with the initial reference low-pass filtered to 60 Å routinely.

Beyond resolving significantly more conformers from the same dataset, AlphaCryo4D also allows us to push the envelope of the achievable resolution of dynamic components and key metastable states due to its advantage in keeping more particles without sacrificing conformational homogeneity. To improve the resolution of state E_D2_, a rigorous particle voting procedure was performed in each cluster on the unmasked energy landscape (Figures 2B and 2C), which improved the overall quality of particle dataset while classifying significantly more particles to this E_D2_ conformational class. Next, we conducted a focused 3D classification by using AlphaCryo4D to calculate the zoomed-in energy landscape of the entire E_D2_ dataset to detect any potential conformational changes within this class. Five clusters near the energy wells of state E_D2_, whose particle numbers were 154,661, 243,799, 198,221, 179,257 and 106,047, were investigated by separate high-resolution 3D refinement and reconstruction. The resulting refined density maps at 3.0-3.5 Å were found to be all in an identical conformation as previously reported ^6^. These five clusters were then merged together to refine a final 3D density map of state E_D2_. Thus, AlphaCryo4D allowed us to obtain a new E_D2_ dataset that appears to retain high conformational homogeneity while including three-fold more particles, which improves its 3D reconstruction from the previously published resolution at 3.2 Å to 2.5 Å, measured by gold-standard Fourier shell correlation (FSC) at 0.143-cutoff. By contrast, in our previous work^6^, we had attempted to include more E_D2_-compatible 3D classes from 3D maximum-likelihood classification in RELION, which resulted in lower resolution at 3.3 Å by doubling the E_D2_ dataset, indicating that including more particles by conventional methods gave rise to higher heterogeneity in the class presumably due to increased misclassification.

### Analysis of doubly and singly capped proteasomes

Previous cryo-EM analyses of the proteasomal states were nearly all focused on the RP-CP subcomplex, by converting double-cap (DC) proteasome particles to pseudo single-cap (SC) proteasome, which was always mixed with true SC particle images, in order to improve the resolution with conventional methods.^6,7,13,14,66,67^ As a result, the cryo-EM density of the CP gate in the other side distal to the RP in the RP-CP subcomplex has been an average of closed and open states with unknown stoichiometry. This has made it impossible to appreciate the difference between the SC and DC proteasomes in the previous studies. To remove this ambiguity and local conformational heterogeneity, we reconstructed 8 density maps of the pure SC proteasome and 36 density maps of the pure DC proteasome including all possible combination of the 8 major RP-CP states by AlphaCryo4D classification. Both CP gates in the 44 states of the pure SC and DC reconstructions are expected to be in a homogeneous conformation. To reconstruct the density maps of the DC proteasome, pseudo-SC particles corresponding to 8 different RP states were voted out from the energy landscape, with the sub-states belonging to the same RP state combined together. Then the particles with one RP-CP were extracted from the particle stacks to refine the RP and CP density maps with RP and CP contour masks applied in the local search step in RELION, respectively. For each reconstruction of the DC proteasome, two RP density maps on the opposite sides of the CP were expanded from 600 × 600 × 600 pixels to 800 × 800 × 800 pixels using RELION and aligned by the density of CP in Chimera before being merged together. All 8 SC and 29 DC states were refined to 3.7-4.7 Å, with 7 DC states limited to 6-9 Å due to their transient nature and extremely low population (Figures S4, S4B, S4E and S4F).

Counting the particle number of the 36 states of the DC proteasome, we could derive the distribution of the DC states, p_*ij*_, and that of RP p_*i*_ only, where *i* and *j* denote the states of two RPs in the same DC proteasome. Based on the value of p_*ij*_ and total particle number, the experimental state distribution of the DC proteasome was plotted for quantitative analysis in two dimensions with respect to the states of two RPs in the same DC proteasome (Figure 1E). If the states of two RPs of the same DC proteasome were independent of each other, the predicted state distribution p∼_*ij*_ of the DC proteasome can be calculated as:

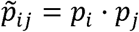

By using the total particle number and experimentally measured p_i_, the 2D state distribution of the DC proteasome with two uncoupled RPs was calculated for further comparison to investigate the conformational entanglement effect of the two RP states (Figure 1F–1H).

### Atomic model building and refinement

To build the initial atomic model of the newly discovered states, we use previously published proteasome structures as a starting model and then manually improved in Coot^68^. For the conformational states at resolutions lower than 5 Å, pseudo-atomic modelling was conducted in the following steps. First, each subunit was fitted as a rigid body, often in UCSF Chimera^69^ as well as in Coot^68^. Then, the local structural domains and secondary structure elements were fitted as a rigid body, with linking loops flexibly fitted and modelled. After the mainchain structures were well fitted, no further fitting of the sidechain rotamers were pursued due to insufficient resolution except for energy minimization to remove unrealistic side-chain clashes. For the conformational states at resolutions higher than 5 Å, the mainchain traces were first fit in Coot. Then, extensive fitting, adjustment and optimization of sidechain rotamers were conducted through local real-space refinement and manual rectification in Coot to the degree commensurate to the map resolution and quality. Atomic model refinement was conducted in Phenix^70^ with its real-space refinement program ^71^. We used both simulated annealing and global energy minimization with secondary structure, rotamer and Ramachandran constraints. Partial rebuilding, model correction and density-fitting improvement in Coot^68^ were iterated after each round of atomic model refinement in Phenix^70^. The improved atomic models were then refined again in Phenix, followed by rebuilding in Coot^68^. The refinement and rebuilding cycle were often repeated for three rounds or until the model quality reached expectation (Table S1). All figures of structures were plotted in Chimera^69^, PyMOL^72^, or ChimeraX^73^. Local resolutions of cryo-EM density maps were evaluated using ResMap^74^ or Bsoft Blocres program^75^. Structural alignment and comparison were performed in both PyMOL and Chimera. Electrostatic surfaces were calculated by APBS plugin^76^ in PyMOL.

Given a large range of variations of local resolution in an expanded number of proteasome states and substates, as well as reconstructions with different stoichiometric ratios of RP versus CP, additional caveats and cautions were practiced in order to avoid misinterpretation and overall-fitting of atomic and pseudo-atomic models. First, when fitting the ubiquitin structure to the low-resolution local density, we consider both the existing homologous structural models as a modelling reference. Specifically, the fitting of diubiquitin to the density on RPN2 in states E_A2.1_, E_A2.2_, and E_B.2_, we took the NMR structure of the yeast diubiquitin-bound Rpn1 T1 site as a reference, because of the high structural homology between RPN1 and RPN2. For the fitting of RPN10-bound density, we also took into account the electrostatic complementarity (Figure S7E-S7H). Second, even within an overall high-resolution cryo-EM map, it often presents certain local densities at a lower local resolution due to poor occupancy of ligands or flexibility of less well-folded segments. For example, for the nucleotide in RPT5 in states E_D0.1_ to E_D1.3_, the high-resolution atomic model of ADP refined from other states like E_C1_ were used to fit these lower-local-resolution densities as a rigid body. However, the B-factors of the fitting atoms of ADP in RPT5 were reported to be more than twice higher than those of other nucleotides, indicating its partial occupancy or unstable binding. Thus, ADP fitting in RPT5 should be regarded tentative for the most or for the purpose of evaluation of nucleotide states rather than a reliable high-resolution atomic modeling. Similarly, the atomic models of these locally low-resolution features were only further adjusted with strict stereochemical constraints when the map resolution and features permit. Third, for the CP gate at the other side opposite to the RP in the RP-CP subcomplex reconstructed from a mixture of true and pseudo-SC proteasomes (converted from the DC proteasomes), their local densities in states E_D0.1_ to E_D2_ appear to be an average of the open and closed states, and thus are weaker in amplitude than the rest of the CP. In the atomic model building of these reconstructions, we chose to model them with the closed CP gate state as long as their density resolution allows atomic modeling. This issue was completely solved when we separated the 3D classes of the SC and DC proteasomes and reconstructed their states respectively, where the atomic modelling of any CP gate no longer suffers from such ambiguities.

### Molecular dynamics simulations analysis

The MD simulations were performed using GROMACS of 2021 version.^77,78^ CHARMM36 all-atom force field was used to build protein topology and the SPC/E water model was added as solvent molecules.^79,80^ The system was constructed in a cubic box at the distances of one nanometer between the solute and the box boundary. Besides, the charge of the ubiquitin-α5 system is neutralized electrically with Na^+^ or Cl^−^ ions, and the system is in the environment with salt solution of 0.154 mol/L NaCl (Normal saline concentration).^81,82^ After the system is assembled, energy minimization, NVT and NPT equilibration are performed to ensure that the system is well-equilibrated at the desired temperature (300 K) and pressure (1 bar).^83^ During this time, the cutoff of short-range electrostatic and van der Waal interactions were both 1 nm, while for long-range Coulomb interactions, they were treated with Particle Mesh Ewald (PME) method.^84^ The final MD production of the system continued up to 100[ns, at the time steps of 2 fs. Since the simulation was conducted with periodic boundary conditions, we recentered the protein and rewrap the molecules within the unit cell to prevent molecules from jumping back and forth across the box boundary after the simulation was done. We analyzed the system by calculating RMSD based on the following formula, which reflects the conformational change of the molecule.

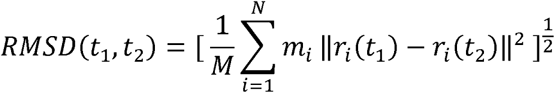

where 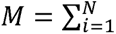 and r_i_ (t) is the position of atom *i* at time t.

### Quantum mechanical calculation

First-principles calculations on the electronic structures of the substrate-proteolytic site complex were conducted as described. In brevity, local coordinates of the substrate-bound CP near the catalytic site at residue Thr1 were taken out from the experimentally fitted atomic models of CP and the substrate, in both E_A_ and E_D2_ conformations. The proteolytic site is the Thr1 residue at the β2 subunit but extended along the chain for another two residues to make the system less finite. Hydrogen atoms are added to complete the residues, and to saturate the system boundaries. The structural relaxations and electronic structure calculation were carried out using the density functional theory (DFT) method with norm-conserving pseudopotentials as implemented in the SIESTA code.^85^ To set up the initial conditions of the finite system, the hydrogen atom in the hydroxy group of Thr1 is manually relocated to the nearby nitrogen atom in the same residue, corresponding to the initial proton transfer in the catalytic mechanism. All hydrogen atoms were subsequently optimized without symmetric restrictions using the conjugate gradient algorithm and a 0.04 eV/Å maximum force convergence criterion while keeping the rest of the system fixed. The generalized gradient approximation (GGA) exchange-correlation density functional PBE^38^ was employed together with a double-zeta plus polarization basis set, and a mesh cutoff of 200 Ry (corresponds to 0.23 Å smallest grid size). The charge density and charge density difference contour maps were plotted with the Siesta Utility programs denchar and Python, respectively.

## Data availability

The atomic coordinates and their corresponding cryo-EM density for all 64 proteasome conformers have been deposited into the RCSB Protein Data Bank (PDB) (www.rcsb.org) and Electron Microscopy Data Bank (EMDB) (www.emdatasource.org). Raw image data have been deposited into the Electron Microscopy Pilot Image Archive (www.ebi.ac.uk/pdbe/emdb/empiar) under the accession numbers EMPIAR-10669, for the cryo-EM dataset of the substrate-engaged human 26S proteasome.

## Acknowledgments

The authors thank A. Goldberg, D. Finley, S. Elsasser, M. Kirschner, Y. Lu, and for constructive discussions. The cryo-EM data were collected at the Cryo-EM Core Facility Platform and Laboratory of Electron Microscopy at Peking University. The data processing was performed in the High-Performance Computing Platform at Peking University. This work was supported by National Key Research and Development Program of China (2023YFF1204400 and 2023YFF1204401), National Natural Science Foundation of China (grant no. 12125401, 12090051, 11774012 and 32471308), Beijing Natural Science Foundation (grant Z180016/Z18J008), and AI for Science (AI4S)-Preferred Program at Peking University Shenzhen Graduate School.

## Author contributions

Y.M. conceived this study. S.Z., Y.D., Z.W. and W.L.W. contributed to the experimental cryo-EM data acquisition. Z.W. and S.Z. analyzed the experimental cryo-EM datasets and refined the density maps. E.C. and W.L.W. conducted the quantum mechanical simulation. Z.W. and Y.M. verified the density maps, built and refined the atomic models and analyzed the data. Y.M. supervised this study and wrote the manuscript with inputs from all authors.

