## Supplementary Information for "Hidden dynamics of ubiquitin-mediated autoregulation of human 26S proteasome during substrate degradation"

**This file includes:**

Figures S1 to S8
Table S1
Captions for Movies S1 to S3

**Other Supplemental Information files for this manuscript include the following:**

Movies S1 to S3

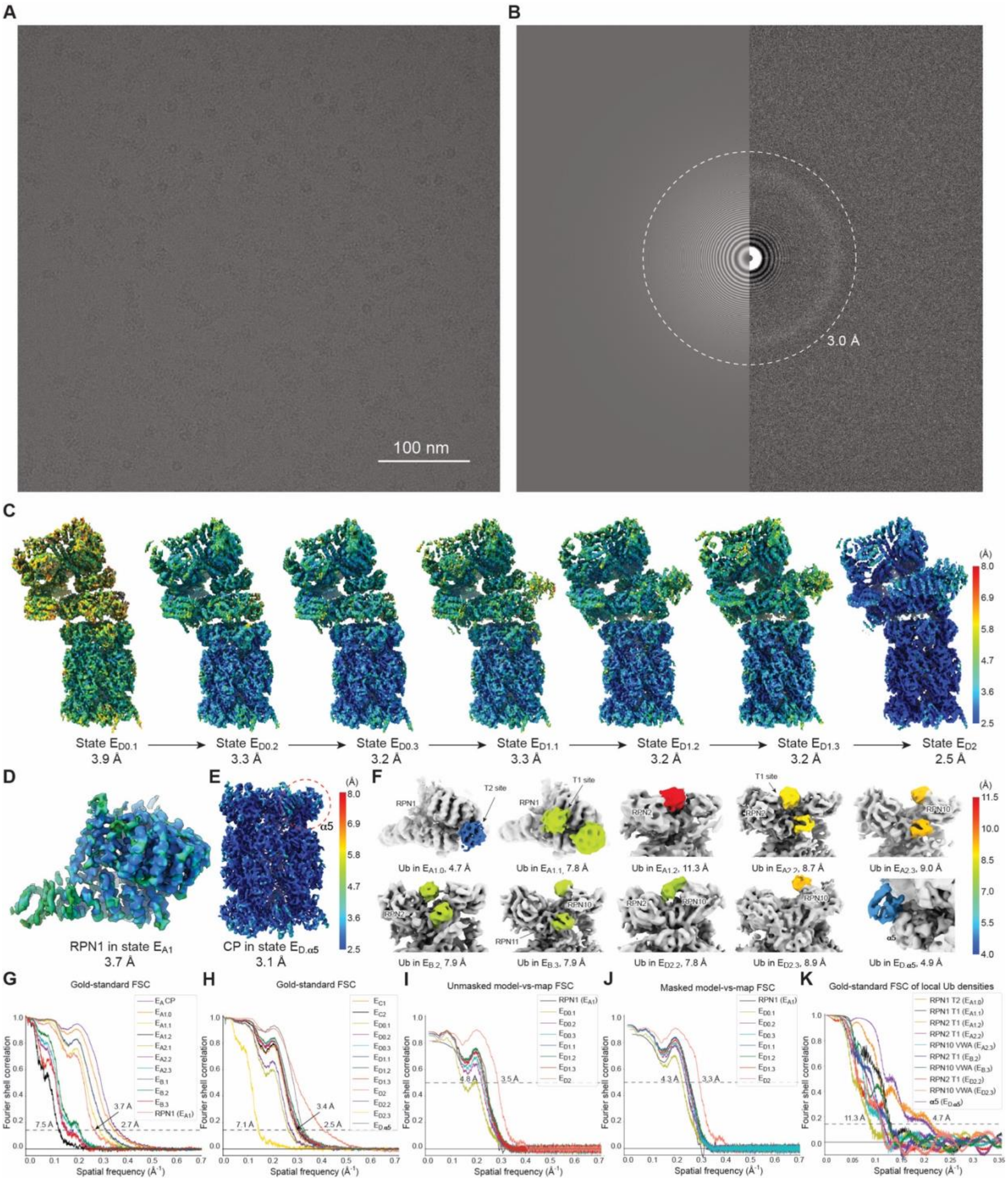

**Figure S1. Cryo-EM reconstructions of novel proteasomal states.**

(**A**) A typical raw micrograph of the substrate-bound human proteasome recorded under the super-resolution counting mode of Gatan K2 Summit direct electron detector. (**B**) The power spectrum of the micrograph shown in panel (**A**), with the 3 Å resolution ring marked by the dashed circle. (**C**) Local resolution measurement of the seven sequential intermediate states during translocation initiation measured by ResMap (Kucukelbir et al., 2014). (**D**) The local resolution measurement of the improved density map of RPN1 in state $E_{A1}$ by ResMap. (**E**) The local resolution measurement of the CP in state $E_{D.\alpha5}$ by ResMap. (**F**) Putative ubiquitin (Ub) densities in ten distinct states colored by the local gold-standard FSC resolutions of the Ub densities measured in panel (**K**), with the rest of densities shown in light grey. The color bar for local resolutions is shown on the right in panels in **C**-**F**, where panels (**D**) and (**E**) share the same resolution color bar. (**G** and **H**) The gold-standard FSC plots of the 20 conformers of the RP-CP subcomplex. (**I** and **J**) The FSC curves calculated between the experimental cryo-EM maps and their corresponding atomic models for newly discovered states or improved states without (**I**) and with (**J**) masking in the calculations. **(K**) The gold-standard FSC curves calculated between the two half maps with local Ub masks in ten distinct states.

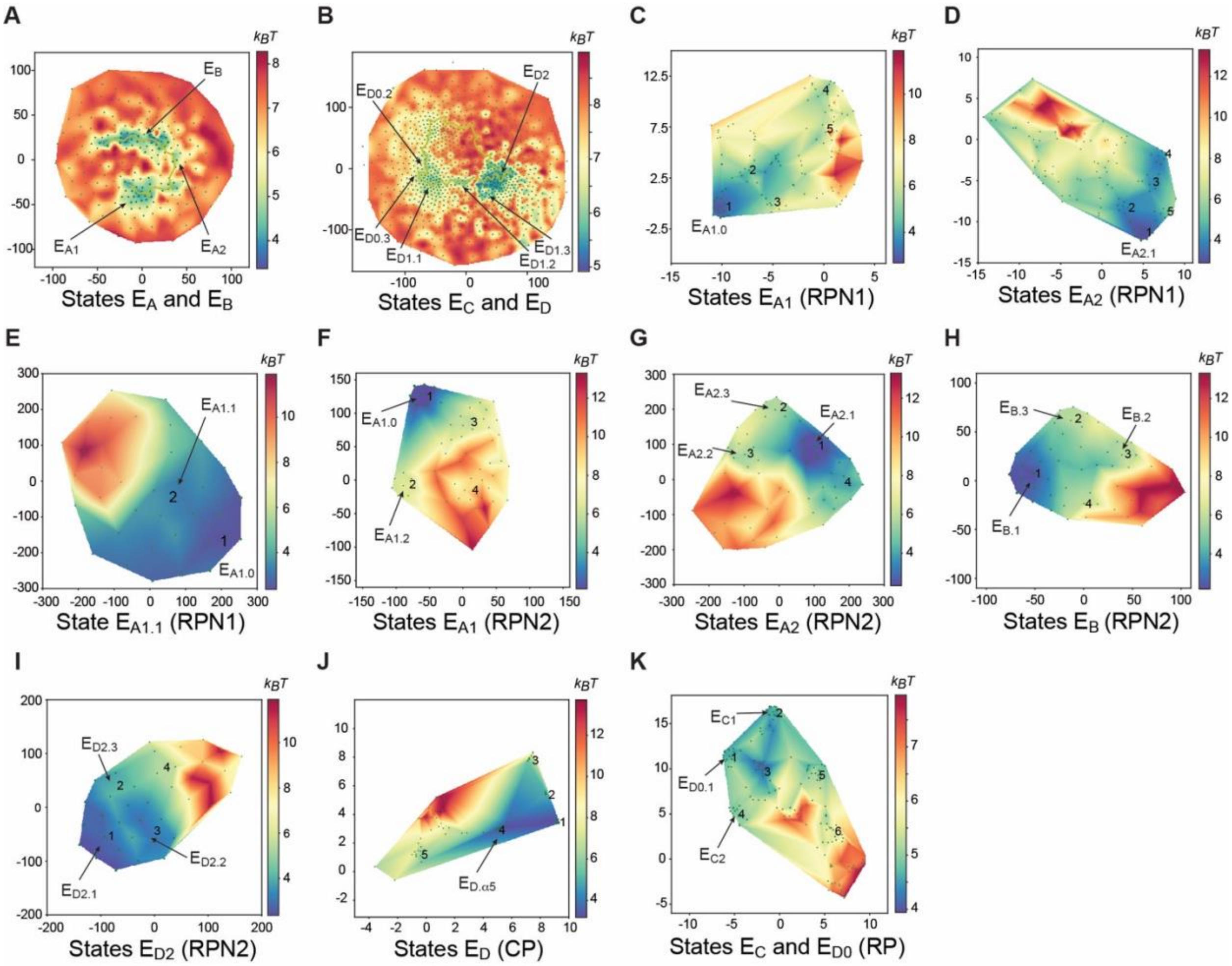

**Figure S2. Energy landscapes of substrate-engaged human proteasome computed to guide in-depth focused 3D classification.**
(**A**-**K**) The zoomed-in energy landscape used for focused 3D classifications using program AlphaCryo4D for finding new states of the substrate-bound proteasome. The zoomed-in local energy landscapes of the substrate-bound human proteasome by AlphaCryo4D are shown, respectively, for state $E_A$ and $E_B$ in (**A**), for states $E_C$ and $E_D$ in (**B**), for states $E_{A1}$ and $E_{A2}$ with 3D mask focusing on RPN1 in (**C**) and (**D**) to visualize the RPN1 T2 site, in (**E**) to visualize the RPN1 T1 site, for states $E_{A1}$, $E_{A2}$, $E_B$ and $E_{D2}$ with 3D masking around RPN2 in (**F**), (**G**), (**H**) and (**I**) to visualize the RPN2 T1 site, for combined states of $E_D$ with masking around CP (**J**) to visualize the ubiquitin-binding sites on the α5-subunit, and for combined states of $E_C$ and $E_{D0}$ with 3D mask focusing on the RP (**K**) to visualize the ATPase motor intermediates during translocation initiation.

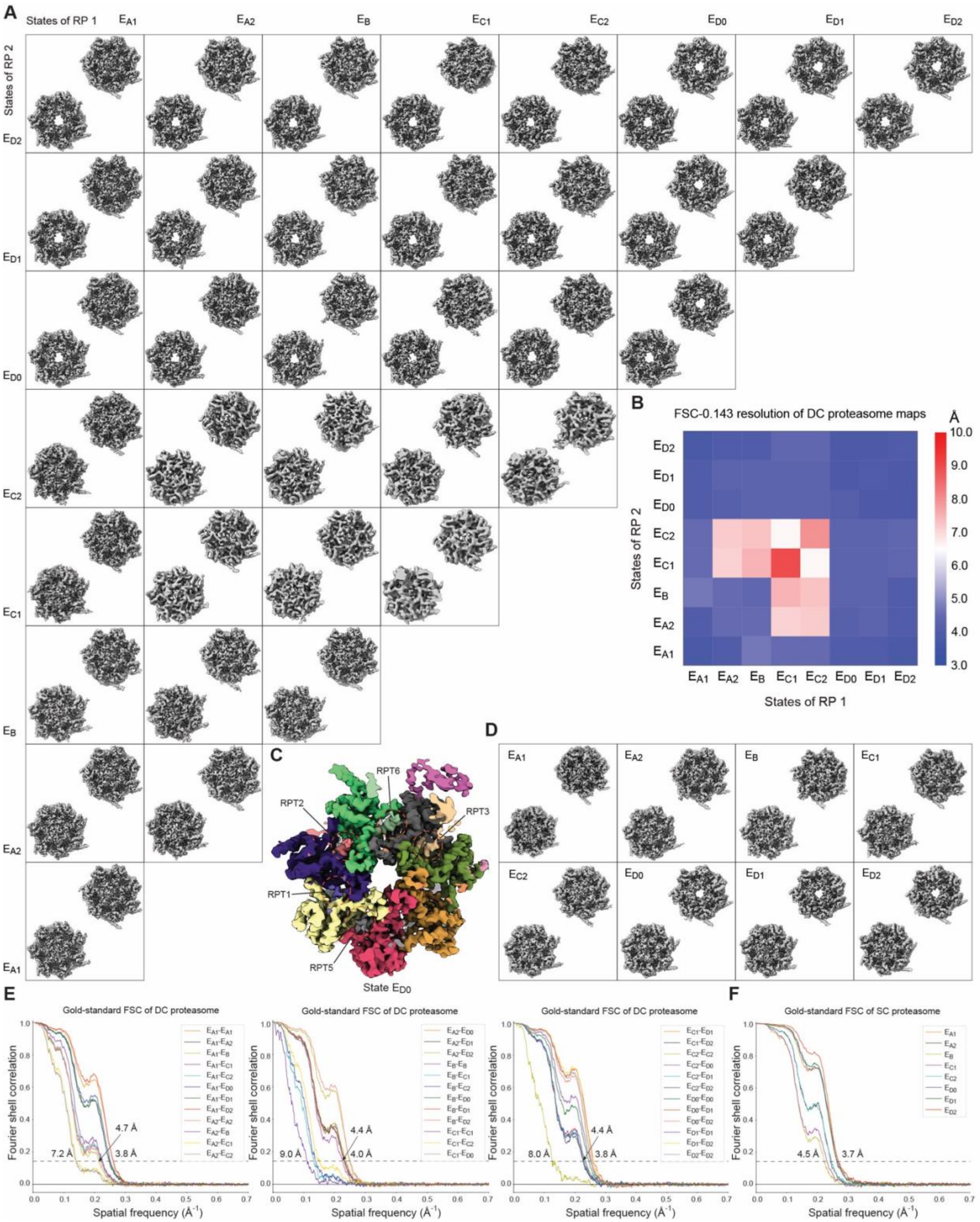

**Figure S3. Cryo-EM analysis of the doubly capped (DC) and singly capped (SC) proteasomes.** (**A**) The CP gate states of the 36 density maps of distinct conformational states of the DC proteasome. All density maps shown are low-pass filtered to their measured resolutions by the gold-standard FSC-0.143 cutoff without amplitude correction of B-factors. (**B**) The FSC-0.143 resolution matrix of the 36 DC proteasomal states. (**C**) The RP-CP interface of state $E_{D0}$, showing five C-terminal tails inserted into the inter-subunit surface pockets of α-ring. (**D**) The CP gate states of the 8 density maps of distinct conformational states of the SC proteasome, with the upper right image in each panel corresponding to the RP-proximal side of the CP gate, and the lower left image to the RP-distal side of the CP gate. (**E**) The gold-standard FSC plots of the 36 conformers of the DC proteasome. (**F**) The gold-standard FSC plots of the 8 conformers of the SC proteasome.

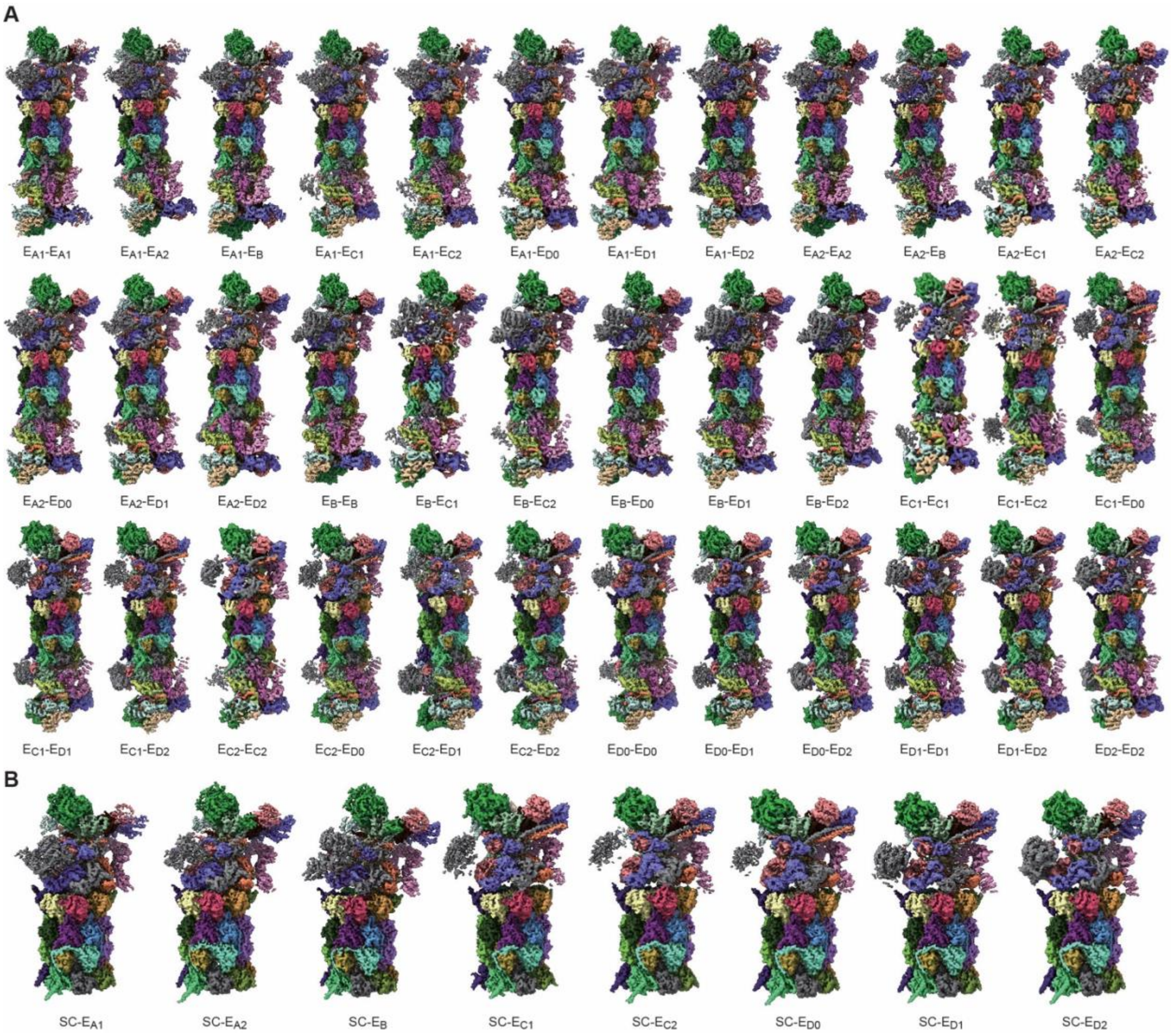

**Figure S4. Cryo-EM reconstructions of the SC and DC proteasomes.**
(**A**) The density maps of 36 distinct conformers of the DC proteasome classified by AlphaCryo4D. (**B**) The density maps of 8 distinct conformers of the SC proteasome. All maps are filtered to their respective gold-standard FSC-0.143 resolutions without amplitude correction of B-factors and are differentially colored by their subunits in ChimeraX (Goddard et al., 2018).

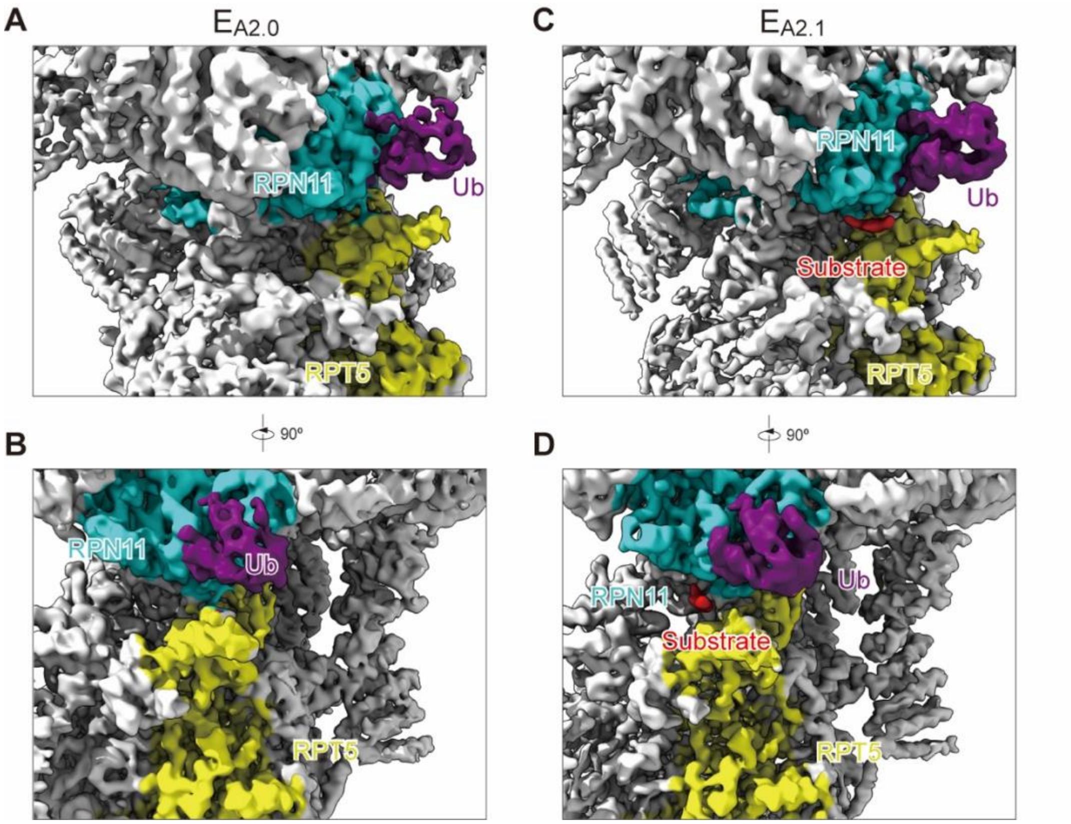

**Figure S5**. **A newly identified intermediate state $E_{A2.1}$ between states $E_{A2}$ and $E_B$.** (**A** and **B**) Close-up view of the ubiquitin interaction with RPN11 in state $E_{A2.0}$ shown in two orthogonal perspectives. The RP conformation in state $E_{A2.0}$ is identical to the previously reported state $E_{A2}$, but the local resolution of the RPN11-bound ubiquitin is improved in state $E_{A2.0}$ compared to that of state $E_{A2}$. (**C** and **D**) Close-up view of the ubiquitin interaction with RPN11 in state $E_{A2.1}$ shown in two orthogonal perspectives, where a short segment of substrate density (highlighted as red) is observed to bind the cleft between RPN11 and RPT4 near the active site of RPN11. While both states $E_{A2.0}$ and $E_{A2.1}$ show very similar AAA-ATPase conformations that are not yet engaged with the substrate, the lid is also rotated by a small angle in state $E_{A2.1}$ relative to state $E_{A2.0}$. The RPT5 N-loop, C-terminal strand of ubiquitin and insert-1 hairpin of RPN11 already form a four-stranded b-sheet in both states $E_{A2.0}$ and $E_{A2.1}$, suggesting that the formation of the four-stranded b-sheet on RPN11-ubiquitin interface proceeds the substrate binding to the RPN11-RPT4 cleft. These features indicate that state $E_{A2.1}$ represent an intermediate between states $E_{A2}$ and $E_B$ that proceeds the substrate insertion into the AAA-ATPase and deubiquitylation.

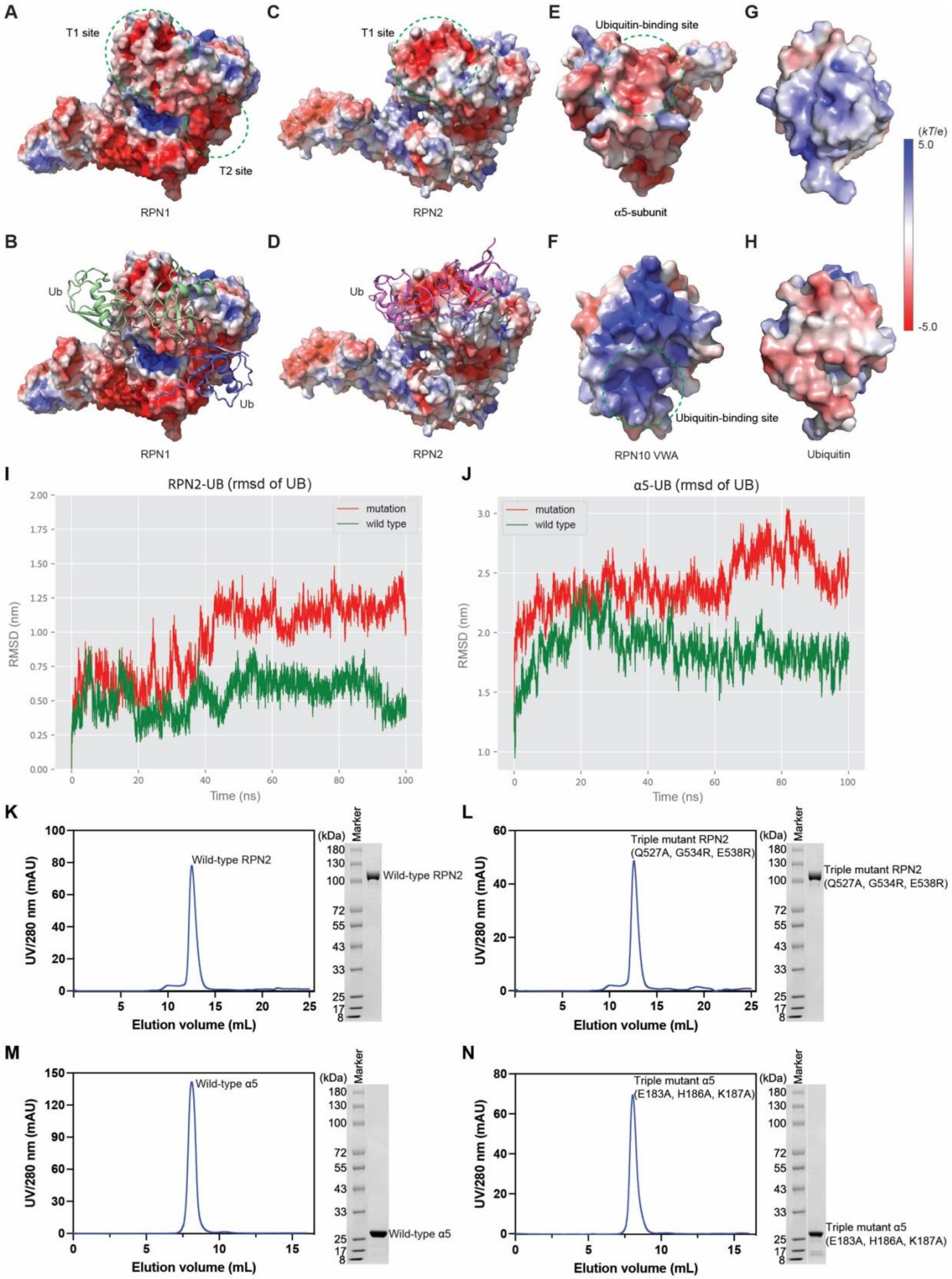

A
T1 site
T2 site
RPN1
B
Ub
Ub
RPN1
C
T1 site
RPN2
D
Ub
RPN2
E
Ubiquitin-binding site
α5-subunit
F
Ubiquitin-binding site
RPN10 VWA
G
H
Ubiquitin
(kT/e)
5.0
-5.0
I
RPN2-UB (rmsd of UB)
mutation
wild type
RMSD (nm)
Time (ns)
J
α5-UB (rmsd of UB)
mutation
wild type
RMSD (nm)
Time (ns)
K
Wild-type RPN2
UV/280 nm (mAU)
Elution volume (mL)
(kDa)
Marker
Wild-type RPN2
L
Triple mutant RPN2
(Q527A, G534R, E538R)
UV/280 nm (mAU)
Elution volume (mL)
(kDa)
Marker
Triple mutant RPN2
(Q527A, G534R, E538R)
M
Wild-type α5
UV/280 nm (mAU)
Elution volume (mL)
(kDa)
Marker
Wild-type α5
N
Triple mutant α5
(E183A, H186A, K187A)
UV/280 nm (mAU)
Elution volume (mL)
(kDa)
Marker
Triple mutant α5
(E183A, H186A, K187A)

**Figure S6. Computational analysis of ubiquitin (Ub) interactions with RPN1, RPN2 and the α5 subunit**

(**A-F**) The electrostatic surfaces of the RPN1, RPN2, RPN10 and α5 subunits show the charge complementarity of the RPN1 T1 and T2 (panels **A**, **B**), RPN2 T1 (panels **C**, **D**) and α5-binding (panel **E**) sites of Ub interactions are all acidic. By contrast, the RPN10 VWA site is basic (panel **F**), suggesting that it may bind the acidic side of Ub. (**G** and **H)** The electrostatic surface of Ub shows one side that is basic (panel **G**) and the other side acidic (panel **H**). (**I** and **J**) The root mean square deviation (RMSD) profile of Ub during 100 ns molecular dynamics simulation-generated trajectory for RPN2-Ub system (**I**) or α5-Ub system (**J**). The mutants of RPN2 or α5 subunit refer to the triple mutant constructs described in the main paper. (**K-N**) FPLC purification of human wild-type RPN2 (left panel, **K**) and triple mutant RPN2 (left panel, **L**) on Superdex 200 10/300 GL column, and FPLC purification of human wild-type α5 (left panel, **M**) and triple mutant α5 (left panel, **N**) on Superdex 75 10/300 GL column. SDS-PAGE and Coomassie blue stain analysis are shown for purified wild-type RPN2 (right panel, **K**), triple mutant RPN2 (right panel, **L**), wild-type α5 (right panel, **M**) and triple mutant α5 (right panel, **N**).

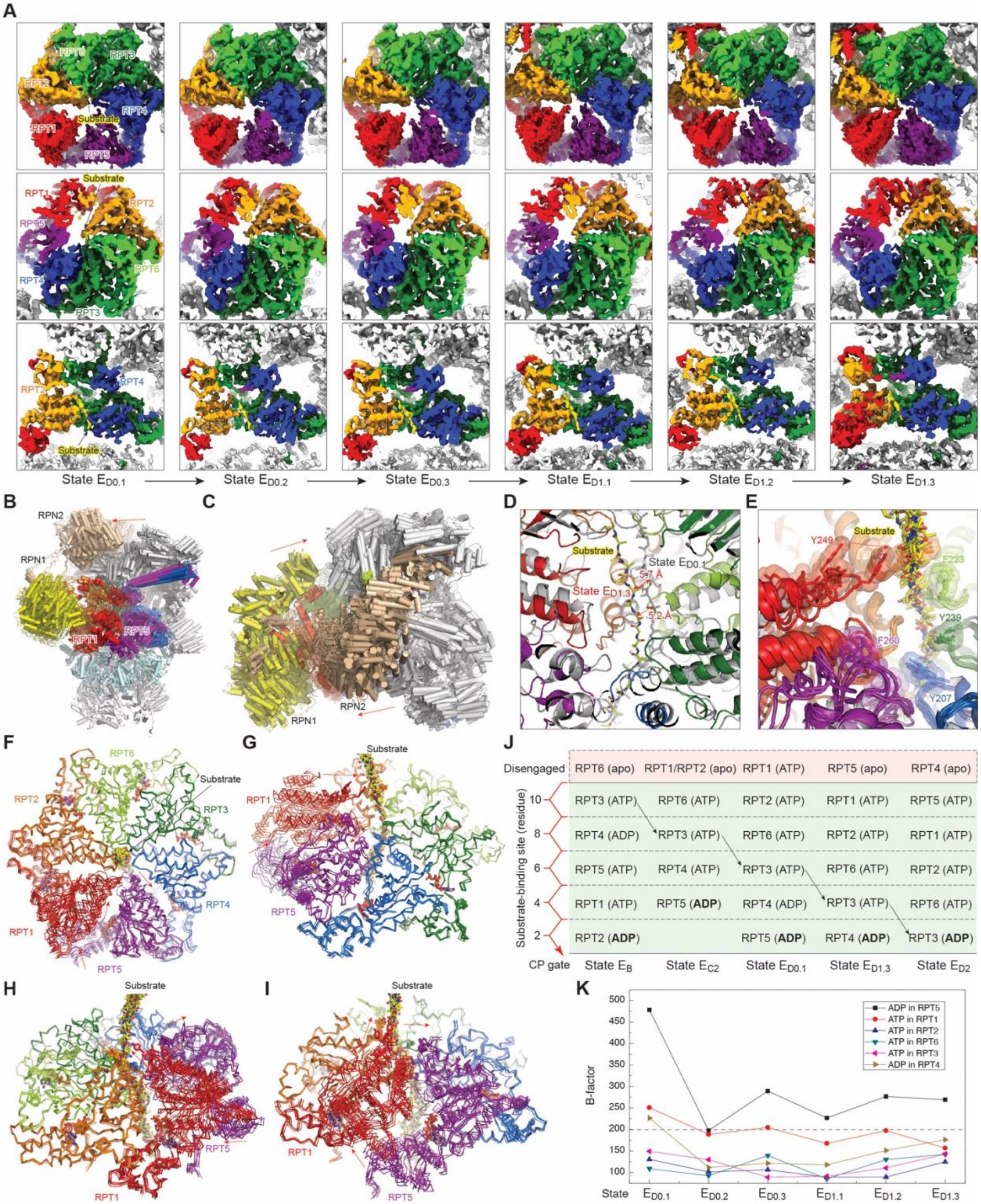

**Figure S7. Cryo-EM structures of six sequential intermediate states between $E_{C2}$ and $E_{D1.3}$.** (**A**) Cryo-EM density maps of the AAA-ATPase motor in six sequential states from $E_{D0.1}$ to $E_{D1.3}$ from three different viewing angles. The substrate densities are highlighted in yellow. (**B**) Superposition of the seven structures in cartoon representations from a lateral perspective, with all structures aligned together based on their CP structures, showing the RP movements relative to the CP. (**C**) Superposition of the six structures in cartoon representations from a top-view perspective, with all structures aligned together based on their CP structures. (**D**) Measurement of substrate movement between states $E_{D0.1}$ and $E_{D1.3}$ relative to the CP, suggesting that the substrate is translated ~5-6 Å during the process of ADP release in RPT5 and of RPT1 re-association with the substrate. (**E**) The closeup view of the pore-1 loop interaction with the substrate with all six states superimposed after aligned together based on the structures of RPT3, RPT4 and RPT6. (**F**-**I**) Superposition of the six sequential states with their RPT3, RPT4 and RPT6 aligned together. Red arrows show the direction of subunit movements in RPT1 and RPT5. (**J**) A diagram illustrating the axial stepping of the substrate-interacting pore-1 loops that is coupled with ATP hydrolysis in the RPT subunits, revised from the previously published results based on the present study (Dong et al., 2019). (**K**) The average B-factor of the nucleotides in six ATPase subunits fitted to the cryo-EM densities of states $E_{D0.1}$ to $E_{D1.3}$ and computed by Phenix in real-space refinement procedure (Adams et al., 2010). The high B-factor of ADP fitted in RPT5 indicates its low occupancy or unstable association and suggests that the ADP in RPT5 undergoes dissociation in this process.

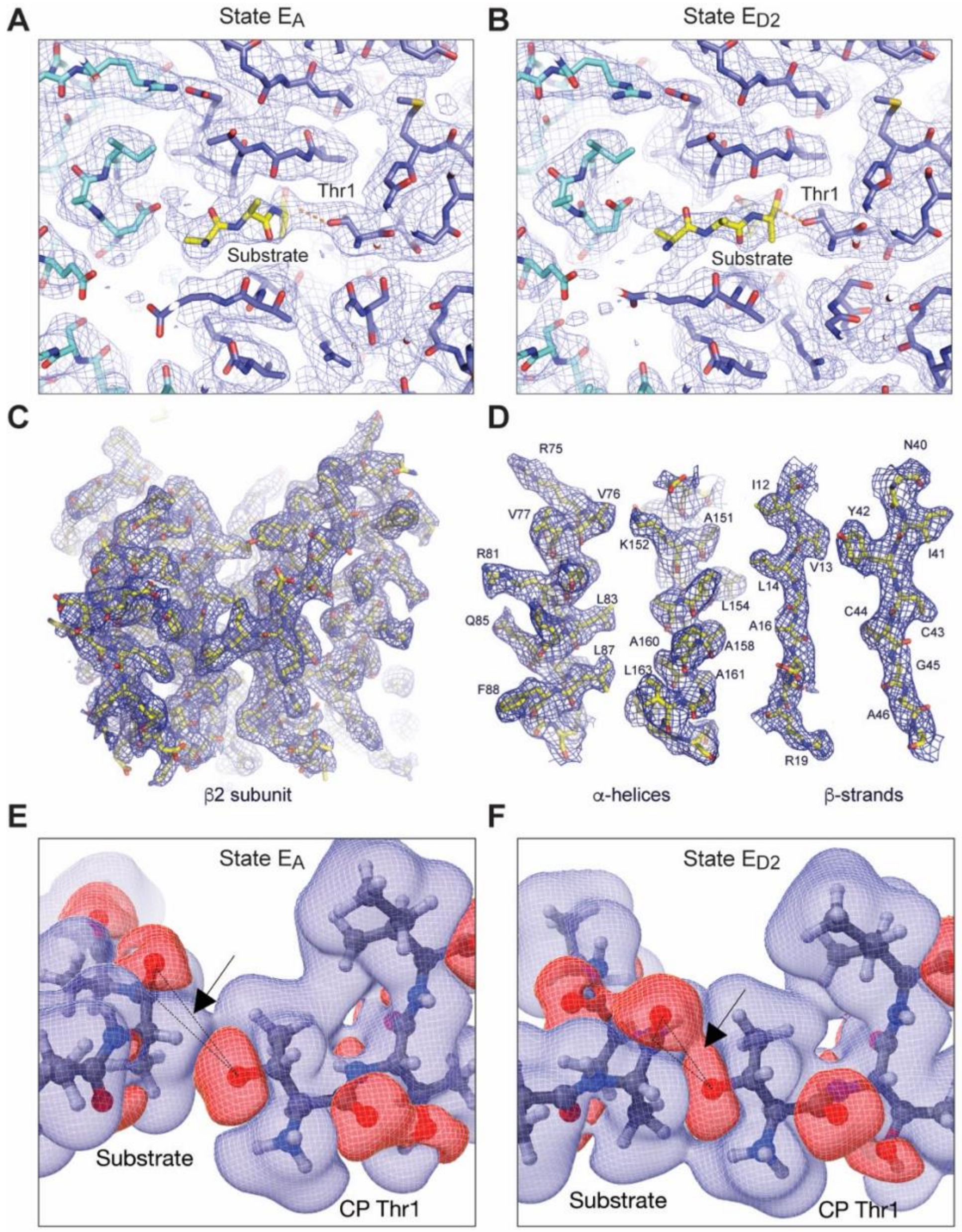

**Figure S8. Substrate interactions with the proteolytic sites in two major CP states.** (**A**) The atomic model of substrate-bound catalytic active site of the β2-subunit superimposed with the cryo-EM density map of state $E_A$ at 2.7 Å. (**B**) The atomic model of substrate-bound catalytic active site of the β2-subunit superimposed with the cryo-EM density map of state $E_{D2}$ at 2.5 Å. (**C**) Closeup view of the cryo-EM density of the β2-subunit in state $E_{D2}$ shown in blue mesh representation superimposed with the atomic model. (**D**) Cryo-EM densities of the typical secondary structure elements in the β2-subunit of state $E_{D2}$ shown as blue mesh representation superimposed with their corresponding atomic models. The residues are labelled. The densities are shown at 4s level and exhibit structural features consistent with 2.5-Å resolution. (**E** and **F**) 3D Charge density difference map showing the interactions between the substrate polypeptide and the residue Thr1 in the β2-subunit of the CP. The red color labels an iso-surface of charge increase while the blue color labels charge decrease, both at a level of 0.04 e/Å$^2$. The charge increase region at the Thr1-Oγ atom is elongated sideways, and the substrate position in the $E_{D2}$ state provides both a proximity and orientation alignment for the pair to interact. The dashed line triangle shows the plane in which the detailed charge difference contours are plotted as shown in Figures 6C and 6D.

### Table S1. Cryo-EM data collection, refinement and validation statistics.

| | $E_{A1.0}$ (EMD-62061) (PDB 9K4J) | $E_{A1.1}$ (EMD-62062) (PDB 9K4K) | $E_{A1.2}$ (EMD-62063) (PDB 9K4L) | $E_{A2.1}$ (EMD-62064) (PDB 9K4M) | $E_{A2.2}$ (EMD-62065) (PDB 9K4O) | $E_{A2.3}$ (EMD-62066) (PDB 9K4P) | $E_{B.1}$ (EMD-62067) (PDB 9K4R) | $E_{B.2}$ (EMD-62068) (PDB 9K4S) |
|---|---|---|---|---|---|---|---|---|
| **Data collection and processing** | | | | | | | | |
| Magnification | 105,000 | 105,000 | 105,000 | 105,000 | 105,000 | 105,000 | 105,000 | 105,000 |
| Voltage (kV) | 300 | 300 | 300 | 300 | 300 | 300 | 300 | 300 |
| Electron exposure (e–/Å²) | 44 | 44 | 44 | 44 | 44 | 44 | 44 | 44 |
| Defocus range (μm) | -0.6 to -3.5 | -0.6 to -3.5 | -0.6 to -3.5 | -0.6 to -3.5 | -0.6 to -3.5 | -0.6 to -3.5 | -0.6 to -3.5 | -0.6 to -3.5 |
| Pixel size (Å) | 0.685 | 0.685 | 0.685 | 0.685 | 0.685 | 0.685 | 0.685 | 0.685 |
| Symmetry imposed | C1 | C1 | C1 | C1 | C1 | C1 | C1 | C1 |
| Initial particle images (no.) | 3,254,352 | 3,254,352 | 3,254,352 | 3,254,352 | 3,254,352 | 3,254,352 | 3,254,352 | 3,254,352 |
| Final particle images (no.) | 192,219 | 43,620 | 2,539 | 147,108 | 5,500 | 5,842 | 173,931 | 9,754 |
| Map resolution (Å) | 2.9 | 4.0 | 7.5 | 3.2 | 6.0 | 4.8 | 3.1 | 4.7 |
| FSC threshold | 0.143 | 0.143 | 0.143 | 0.143 | 0.143 | 0.143 | 0.143 | 0.143 |
| Map resolution range (Å) | 2.5-8.0 | 3.5-8.0 | 7.0-20.0 | 2.5-8.0 | 5.5-20.0 | 4.0-20.0 | 2.5-8.0 | 4.5-20.0 |
| **Refinement** | | | | | | | | |
| Initial model used | 6MSB | | | 6MSD | | | 6MSE | |
| Model resolution (Å) | 3.3 | 4.9 | | 3.5 | | | 3.7 | |
| FSC threshold | 0.5 | 0.5 | | 0.5 | | | 0.5 | |
| Model resolution range (Å) | 2.5-8.0 | 3.5-8.0 | | 2.5-8.0 | | | 2.5-8.0 | |
| Map sharpening *B* factor (Å²) | -30 | -28 | 0 | -30 | 0 | 0 | -30 | 0 |
| **Model composition** | | | | | | | | |
| Non-hydrogen atoms | 106622 | 107223 | 106572 | 105930 | 107132 | 106531 | 106266 | 107468 |
| Protein residues | 13591 | 13667 | 13591 | 13468 | 13620 | 13544 | 13513 | 13665 |
| Ligands | 12 | 12 | 12 | 12 | 12 | 12 | 9 | 9 |
| *B* factors (Å²) | | | | | | | | |
| Protein | 122.75 | 254.54 | 282.27 | 125.82 | 193.35 | 207.63 | 109.81 | 176.30 |
| Ligand | 66.07 | 218.97 | 139.21 | 123.91 | 131.42 | 176.78 | 115.30 | 153.31 |
| **R.m.s. deviations** | | | | | | | | |
| Bond lengths (Å) | 0.005 | 0.005 | 0.005 | 0.005 | 0.005 | 0.005 | 0.005 | 0.006 |
| Bond angles (°) | 1.097 | 1.151 | 1.174 | 1.116 | 1.142 | 1.127 | 1.127 | 1.230 |
| **Validation** | | | | | | | | |
| MolProbity score | 1.63 | 1.78 | 1.88 | 1.66 | 1.90 | 1.84 | 1.67 | 1.91 |
| Clashscore | 4.80 | 6.98 | 8.84 | 5.26 | 9.33 | 7.78 | 4.91 | 9.17 |
| Poor rotamers (%) | 0.26 | 0.21 | 0.29 | 0.21 | 0.37 | 0.36 | 0.31 | 0.57 |
| **Ramachandran plot** | | | | | | | | |
| Favored (%) | 94.46 | 94.09 | 93.89 | 94.53 | 94.04 | 93.84 | 93.85 | 93.69 |
| Allowed (%) | 5.39 | 5.80 | 5.99 | 5.35 | 5.76 | 5.99 | 5.89 | 6.05 |
| Disallowed (%) | 0.15 | 0.10 | 0.12 | 0.12 | 0.21 | 0.17 | 0.26 | 0.26 |

### Cryo-EM data collection, refinement and validation statistics (continued)

| | $E_{B.3}$ (EMD-62069) (PDB 9K4T) | $E_{C1}$ (EMD-62070) (PDB 9K4U) | $E_{C2}$ (EMD-62071) (PDB 9K4V) | $E_{D0.1}$ (EMD-62072) (PDB 9K4W) | $E_{D0.2}$ (EMD-62073) (PDB 9K4X) | $E_{D0.3}$ (EMD-62074) (PDB 9K4Y) | $E_{D1.1}$ (EMD-60675) (PDB 9K4Z) | $E_{D1.2}$ (EMD-62076) (PDB 9K50) |
|---|---|---|---|---|---|---|---|---|
| **Data collection and processing** | | | | | | | | |
| Magnification | 105,000 | 105,000 | 105,000 | 105,000 | 105,000 | 105,000 | 105,000 | 105,000 |
| Voltage (kV) | 300 | 300 | 300 | 300 | 300 | 300 | 300 | 300 |
| Electron exposure (e–/Å$^2$) | 44 | 44 | 44 | 44 | 44 | 44 | 44 | 44 |
| Defocus range (μm) | -0.6 to -3.5 | -0.6 to -3.5 | -0.6 to -3.5 | -0.6 to -3.5 | -0.6 to -3.5 | -0.6 to -3.5 | -0.6 to -3.5 | -0.6 to -3.5 |
| Pixel size (Å) | 0.685 | 0.685 | 0.685 | 0.685 | 0.685 | 0.685 | 0.685 | 0.685 |
| Symmetry imposed | C1 | C1 | C1 | C1 | C1 | C1 | C1 | C1 |
| Initial particle images (no.) | 3,254,352 | 3,254,352 | 3,254,352 | 3,254,352 | 3,254,352 | 3,254,352 | 3,254,352 | 3,254,352 |
| Final particle images (no.) | 5,117 | 68,506 | 77,545 | 25,755 | 105,081 | 169,522 | 92,777 | 175,114 |
| Map resolution (Å) | 6.1 | 3.4 | 3.3 | 3.9 | 3.3 | 3.2 | 3.3 | 3.2 |
| FSC threshold | 0.143 | 0.143 | 0.143 | 0.143 | 0.143 | 0.143 | 0.143 | 0.143 |
| Map resolution range (Å) | 5.5-20.0 | 2.5-8.0 | 2.5-8.0 | 3.0-8.0 | 2.5-8.0 | 2.5-8.0 | 2.5-8.0 | 2.5-8.0 |
| **Refinement** | | | | | | | | |
| Initial model used | | 6MSG | 6MSH | | | 6MSH | | |
| Model resolution (Å) | | 3.9 | 3.8 | 4.3 | 4.1 | 4.0 | 3.9 | 3.8 |
| FSC threshold | | 0.5 | 0.5 | 0.5 | 0.5 | 0.5 | 0.5 | 0.5 |
| Model resolution range (Å) | | 3.0-8.0 | 3.0-8.0 | 3.0-8.0 | 3.0-8.0 | 3.0-8.0 | 3.0-8.0 | 3.0-8.0 |
| Map sharpening *B* factor (Å$^2$) | 0 | -30 | -30 | -20 | -20 | -20 | -20 | -30 |
| **Model composition** | | | | | | | | |
| Non-hydrogen atoms | 106867 | 106690 | 106109 | 106334 | 106334 | 106334 | 106334 | 106307 |
| Protein residues | 13589 | 13563 | 13499 | 13513 | 13513 | 13513 | 13513 | 13513 |
| Ligands | 9 | 9 | 8 | 11 | 11 | 11 | 11 | 10 |
| *B* factors (Å$^2$) | | | | | | | | |
| Protein | 182.15 | 137.62 | 174.53 | 176.91 | 164.70 | 135.95 | 141.82 | 122.56 |
| Ligand | 150.20 | 136.45 | 161.26 | 209.99 | 182.04 | 159.32 | 156.12 | 140.27 |
| **R.m.s. deviations** | | | | | | | | |
| Bond lengths (Å) | 0.005 | 0.006 | 0.005 | 0.006 | 0.005 | 0.006 | 0.005 | 0.005 |
| Bond angles (°) | 1.150 | 1.151 | 1.116 | 1.202 | 1.132 | 1.181 | 1.222 | 1.171 |
| **Validation** | | | | | | | | |
| MolProbity score | 1.90 | 1.78 | 1.71 | 1.86 | 1.75 | 1.79 | 1.77 | 1.76 |
| Clashscore | 9.35 | 6.17 | 5.59 | 7.17 | 6.09 | 6.26 | 6.39 | 6.09 |
| Poor rotamers (%) | 0.57 | 0.39 | 0.31 | 0.56 | 0.46 | 0.32 | 0.52 | 0.42 |
| **Ramachandran plot** | | | | | | | | |
| Favored (%) | 93.99 | 93.10 | 93.89 | 92.52 | 93.80 | 92.97 | 93.66 | 93.51 |
| Allowed (%) | 5.81 | 6.68 | 5.93 | 7.25 | 6.02 | 6.81 | 6.00 | 6.19 |
| Disallowed (%) | 0.21 | 0.22 | 0.18 | 0.22 | 0.19 | 0.22 | 0.34 | 0.29 |

### Cryo-EM data collection, refinement and validation statistics (continued)

| | $E_{D1.3}$ (EMD-62077) (PDB 9K51) | $E_{D2.1}$ (EMD-62079) (PDB 9K53) | $E_{D2.2}$ (EMD-62080) (PDB 9K54) | $E_{D2.3}$ (EMD-62081) (PDB 9K55) | $E_A$ CP (EMD-24254) (PDB 9K4J) | $E_{A1}$ RPN1 (PDB 9K4J) | $E_{D.\alpha5}$ CP (PDB 9K54) |
|---|---|---|---|---|---|---|---|
| **Data collection and processing** | | | | | | | |
| Magnification | 105,000 | 105,000 | 105,000 | 105,000 | 105,000 | 105,000 | 105,000 |
| Voltage (kV) | 300 | 300 | 300 | 300 | 300 | 300 | 300 |
| Electron exposure (e–/Å$^2$) | 44 | 44 | 44 | 44 | 44 | 44 | 44 |
| Defocus range (μm) | -0.6 to -3.5 | -0.6 to -3.5 | -0.6 to -3.5 | -0.6 to -3.5 | -0.6 to -3.5 | -0.6 to -3.5 | -0.6 to -3.5 |
| Pixel size (Å) | 0.685 | 0.685 | 0.685 | 0.685 | 0.685 | 1.37 | 0.685 |
| Symmetry imposed | C1 | C1 | C1 | C1 | C1 | C1 | C1 |
| Initial particle images (no.) | 3,254,352 | 3,254,352 | 3,254,352 | 3,254,352 | 3,254,352 | 3,254,352 | 3,254,352 |
| Final particle images (no.) | 146,506 | 856,683 | 24,674 | 6,192 | 339,327 | 128,161 | 131,570 |
| Map resolution (Å) | 3.2 | 2.5 | 3.7 | 7.0 | 2.7 | 3.7 | 3.1 |
| FSC threshold | 0.143 | 0.143 | 0.143 | 0.143 | 0.143 | 0.143 | 0.143 |
| Map resolution range (Å) | 2.5-8.0 | 2.5-4.8 | 3.0-8.0 | 6.0-20.0 | 2.5-8.0 | 3.0-8.0 | 2.5-10.0 |
| **Refinement** | | | | | | | |
| Initial model used | 6MSJ | 6MSK | | | 6MSB | | |
| Model resolution (Å) | 3.9 | 3.2 | 4.5 | | 3.2 | 3.8 | 3.6 |
| FSC threshold | 0.5 | 0.5 | 0.5 | | 0.5 | 0.5 | 0.5 |
| Model resolution range (Å) | 3.0-8.0 | 2.5-5.8 | 3.5-8.0 | | 2.5-8.0 | 3.5-8.0 | 2.9-5.8 |
| Map sharpening *B* factor (Å$^2$) | -30 | -20 | -40 | 0 | -25 | -100 | -25 |
| **Model composition** | | | | | | | |
| Non-hydrogen atoms | 106307 | 106358 | 108712 | 108111 | 63654 | 7717 | 107626 |
| Protein residues | 13513 | 13513 | 13817 | 13741 | 8203 | 994 | 13724 |
| Ligands | 10 | 11 | 11 | 11 | 11 | | 11 |
| *B* factors (Å$^2$) | | | | | | | |
| Protein | 121.05 | 107.76 | 149.76 | 220.56 | 131.13 | 141.87 | 162.57 |
| Ligand | 145.58 | 113.51 | 166.78 | 211.86 | 125.87 | | 135.00 |
| **R.m.s. deviations** | | | | | | | |
| Bond lengths (Å) | 0.006 | 0.004 | 0.005 | 0.005 | 0.008 | 0.005 | 0.007 |
| Bond angles (°) | 1.171 | 1.072 | 1.142 | 1.161 | 0.908 | 0.837 | 1.088 |
| **Validation** | | | | | | | |
| MolProbity score | 1.80 | 2.01 | 1.72 | 1.95 | 1.49 | 3.41 | 1.88 |
| Clashscore | 6.51 | 9.38 | 5.86 | 9.76 | 2.72 | 24.57 | 6.09 |
| Poor rotamers (%) | 0.46 | 1.42 | 0.30 | 0.43 | 0.47 | 12.01 | 0.73 |
| **Ramachandran plot** | | | | | | | |
| Favored (%) | 93.16 | 94.03 | 94.01 | 93.28 | 93.3 | 81.98 | 90.17 |
| Allowed (%) | 6.53 | 5.89 | 5.75 | 6.43 | 6.52 | 17.51 | 9.4 |
| Disallowed (%) | 0.31 | 0.08 | 0.24 | 0.29 | 0.18 | 0.51 | 0.43 |

### Cryo-EM data collection, refinement and validation statistics (continued)

| | SC-$E_{A1}$ (EMD-62082) (PDB 9K56) | SC-$E_{A2}$ (EMD-62083) (PDB 9K57) | SC-$E_{B}$ (EMD-62084) (PDB 9K58) | SC-$E_{C1}$ (EMD-62085) (PDB 9K59) | SC-$E_{C2}$ (EMD-62086) (PDB 9K5A) | SC-$E_{D0}$ (EMD-62087) (PDB 9K5B) | SC-$E_{D1}$ (EMD-62088) (PDB 9K5C) | SC-$E_{D2}$ (EMD-62089) (PDB 9K5D) |
|---|---|---|---|---|---|---|---|---|
| **Data collection and processing** | | | | | | | | |
| Magnification | 105,000 | 105,000 | 105,000 | 105,000 | 105,000 | 105,000 | 105,000 | 105,000 |
| Voltage (kV) | 300 | 300 | 300 | 300 | 300 | 300 | 300 | 300 |
| Electron exposure (e–/Å$^2$) | 44 | 44 | 44 | 44 | 44 | 44 | 44 | 44 |
| Defocus range (μm) | -0.6 to -3.5 | -0.6 to -3.5 | -0.6 to -3.5 | -0.6 to -3.5 | -0.6 to -3.5 | -0.6 to -3.5 | -0.6 to -3.5 | -0.6 to -3.5 |
| Pixel size (Å) | 0.685 | 0.685 | 0.685 | 0.685 | 0.685 | 0.685 | 0.685 | 0.685 |
| Symmetry imposed | C1 | C1 | C1 | C1 | C1 | C1 | C1 | C1 |
| Initial particle images (no.) | 3,254,352 | 3,254,352 | 3,254,352 | 3,254,352 | 3,254,352 | 3,254,352 | 3,254,352 | 3,254,352 |
| Final particle images (no.) | 24,241 | 14,171 | 6,862 | 9,319 | 15,828 | 42,969 | 60,157 | 113,188 |
| Map resolution (Å) | 3.8 | 4.1 | 4.5 | 4.3 | 4.1 | 3.8 | 3.8 | 3.7 |
| FSC threshold | 0.143 | 0.143 | 0.143 | 0.143 | 0.143 | 0.143 | 0.143 | 0.143 |
| Map resolution range (Å) | 3.0-8.0 | 3.0-8.0 | 3.5-8.0 | 3.3-8.0 | 3.0-8.0 | 3.0-8.0 | 3.0-8.0 | 3.0-8.0 |
| **Refinement** | | | | | | | | |
| Initial model used | | | | | | | | |
| Model resolution (Å) | 4.0 | 4.5 | 4.8 | 4.5 | 4.3 | 4.1 | 4.1 | 4.0 |
| FSC threshold | 0.5 | 0.5 | 0.5 | 0.5 | 0.5 | 0.5 | 0.5 | 0.5 |
| Model resolution range (Å) | 3.0-8.0 | 3.8-8.0 | 4.1-8.0 | 3.7-8.0 | 3.3-8.0 | 3.0-8.0 | 3.0-8.0 | 3.0-8.0 |
| Map sharpening *B* factor (Å$^2$) | 0 | 0 | 0 | 0 | 0 | 0 | 0 | 0 |
| **Model composition** | | | | | | | | |
| Non-hydrogen atoms | 105968 | 105927 | 106247 | 106687 | 106134 | 106328 | 106301 | 106302 |
| Protein residues | 13515 | 13468 | 13511 | 13563 | 13499 | 13513 | 13513 | 13513 |
| Ligands | 12 | 12 | 9 | 9 | 8 | 11 | 10 | 11 |
| *B* factors (Å$^2$) | | | | | | | | |
| Protein | 169.04 | 169.31 | 181.26 | 188.47 | 166.11 | 162.08 | 159.13 | 174.11 |
| Ligand | 120.09 | 149.79 | 171.94 | 194.46 | 189.60 | 190.39 | 184.14 | 179.03 |
| **R.m.s. deviations** | | | | | | | | |
| Bond lengths (Å) | 0.005 | 0.006 | 0.005 | 0.005 | 0.005 | 0.005 | 0.005 | 0.006 |
| Bond angles (°) | 1.125 | 1.170 | 1.157 | 1.139 | 1.129 | 1.159 | 1.151 | 1.161 |
| **Validation** | | | | | | | | |
| MolProbity score | 1.64 | 1.74 | 1.75 | 1.77 | 1.70 | 1.77 | 1.73 | 1.77 |
| Clashscore | 5.24 | 6.11 | 6.21 | 6.16 | 5.37 | 6.12 | 5.84 | 6.22 |
| Poor rotamers (%) | 0.27 | 0.24 | 0.43 | 0.43 | 0.26 | 0.51 | 0.50 | 0.52 |
| **Ramachandran plot** | | | | | | | | |
| Favored (%) | 94.75 | 93.89 | 93.94 | 93.37 | 93.84 | 93.41 | 93.73 | 93.40 |
| Allowed (%) | 5.08 | 5.93 | 5.85 | 6.40 | 5.99 | 6.37 | 5.99 | 6.38 |
| Disallowed (%) | 0.16 | 0.19 | 0.21 | 0.22 | 0.17 | 0.22 | 0.28 | 0.22 |

### Cryo-EM data collection, refinement and validation statistics (continued)

| | $E_{A1}$-$E_{A1}$ (EMD-62090) (PDB 9K5E) | $E_{A1}$-$E_{A2}$ (EMD-62091) (PDB 9K5F) | $E_{A1}$-$E_{B}$ (EMD-62092) (PDB 9K5G) | $E_{A1}$-$E_{C1}$ (EMD-62093) (PDB 9K5I) | $E_{A1}$-$E_{C2}$ (EMD-62094) (PDB 9K5J) | $E_{A1}$-$E_{D0}$ (EMD-62095) (PDB 9K5K) | $E_{A1}$-$E_{D1}$ (EMD-62096) (PDB 9K5L) | $E_{A1}$-$E_{D2}$ (EMD-62097) (PDB 9K5M) |
|---|---|---|---|---|---|---|---|---|
| **Data collection and processing** | | | | | | | | |
| Magnification | 105,000 | 105,000 | 105,000 | 105,000 | 105,000 | 105,000 | 105,000 | 105,000 |
| Voltage (kV) | 300 | 300 | 300 | 300 | 300 | 300 | 300 | 300 |
| Electron exposure (e–/Å$^2$) | 44 | 44 | 44 | 44 | 44 | 44 | 44 | 44 |
| Defocus range (μm) | -0.6 to -3.5 | -0.6 to -3.5 | -0.6 to -3.5 | -0.6 to -3.5 | -0.6 to -3.5 | -0.6 to -3.5 | -0.6 to -3.5 | -0.6 to -3.5 |
| Pixel size (Å) | 0.685 | 0.685 | 0.685 | 0.685 | 0.685 | 0.685 | 0.685 | 0.685 |
| Symmetry imposed | C1 | C1 | C1 | C1 | C1 | C1 | C1 | C1 |
| Initial particle images (no.) | 3,254,352 | 3,254,352 | 3,254,352 | 3,254,352 | 3,254,352 | 3,254,352 | 3,254,352 | 3,254,352 |
| Final particle images (no.) | 14,284 | 9,356 | 3,752 | 3,378 | 3,672 | 11,869 | 12,521 | 29,376 |
| Map resolution (Å) | 3.8 | 4.0 | 4.7 | 4.5 | 4.5 | 4.0 | 4.0 | 3.8 |
| FSC threshold | 0.143 | 0.143 | 0.143 | 0.143 | 0.143 | 0.143 | 0.143 | 0.143 |
| Map resolution range (Å) | 3.5-8.0 | 3.8-8.0 | 4.5-15.0 | 4.0-15.0 | 4.0-15.0 | 3.7-8.0 | 3.7-8.0 | 3.6-8.0 |
| **Refinement** | | | | | | | | |
| Initial model used | | | | | | | | |
| Model resolution (Å) | 4.1 | 4.3 | | | | 4.2 | 4.3 | 4.1 |
| FSC threshold | 0.5 | 0.5 | | | | 0.5 | 0.5 | 0.5 |
| Model resolution range (Å) | 3.6-8.0 | 3.8-8.0 | | | | 3.7-8.0 | 3.8-8.0 | 3.6-8.0 |
| Map sharpening *B* factor (Å$^2$) | 0 | 0 | 0 | 0 | 0 | 0 | 0 | 0 |
| **Model composition** | | | | | | | | |
| Non-hydrogen atoms | 164552 | 162661 | 163030 | 163471 | 162911 | 163758 | 163130 | 163131 |
| Protein residues | 20918 | 20642 | 20686 | 20738 | 20674 | 20764 | 20688 | 20688 |
| Ligands | 24 | 24 | 21 | 21 | 20 | 23 | 22 | 23 |
| *B* factors (Å$^2$) | | | | | | | | |
| Protein | 135.48 | 141.81 | 122.56 | 122.59 | 149.65 | 140.41 | 139.96 | 142.12 |
| Ligand | 84.30 | 111.92 | 116.49 | 146.35 | 139.44 | 192.51 | 154.44 | 136.97 |
| **R.m.s. deviations** | | | | | | | | |
| Bond lengths (Å) | 0.004 | 0.004 | 0.005 | 0.004 | 0.004 | 0.004 | 0.005 | 0.004 |
| Bond angles (°) | 1.089 | 1.097 | 1.138 | 1.115 | 1.108 | 1.118 | 1.137 | 1.117 |
| **Validation** | | | | | | | | |
| MolProbity score | 1.63 | 1.64 | 1.74 | 1.70 | 1.67 | 1.68 | 1.77 | 1.71 |
| Clashscore | 4.85 | 5.11 | 5.84 | 5.47 | 5.19 | 5.22 | 6.20 | 5.36 |
| Poor rotamers (%) | 0.21 | 0.27 | 0.24 | 0.26 | 0.26 | 0.29 | 0.41 | 0.31 |
| **Ramachandran plot** | | | | | | | | |
| Favored (%) | 94.50 | 94.69 | 93.72 | 93.89 | 94.20 | 94.04 | 93.35 | 93.62 |
| Allowed (%) | 5.38 | 5.14 | 6.13 | 5.93 | 5.66 | 5.74 | 6.38 | 6.20 |
| Disallowed (%) | 0.12 | 0.17 | 0.16 | 0.18 | 0.14 | 0.22 | 0.28 | 0.18 |

### Cryo-EM data collection, refinement and validation statistics (continued)

| | $E_{A2}$-$E_{A2}$ (EMD-62098) (PDB 9K5N) | $E_{A2}$-$E_{B}$ (EMD-62099) (PDB 9K5O) | $E_{A2}$-$E_{C1}$ (EMD-62100) (PDB 9K5Q) | $E_{A2}$-$E_{C2}$ (EMD-62101) (PDB 9K5R) | $E_{A2}$-$E_{D0}$ (EMD-62102) (PDB 9K5S) | $E_{A2}$-$E_{D1}$ (EMD-62103) (PDB 9K5T) | $E_{A2}$-$E_{D2}$ (EMD-62104) (PDB 9K5U) | $E_{B}$-$E_{B}$ (EMD-62105) (PDB 9K5V) |
|---|---|---|---|---|---|---|---|---|
| **Data collection and processing** | | | | | | | | |
| Magnification | 105,000 | 105,000 | 105,000 | 105,000 | 105,000 | 105,000 | 105,000 | 105,000 |
| Voltage (kV) | 300 | 300 | 300 | 300 | 300 | 300 | 300 | 300 |
| Electron exposure (e–/Å$^2$) | 44 | 44 | 44 | 44 | 44 | 44 | 44 | 44 |
| Defocus range (μm) | -0.6 to -3.5 | -0.6 to -3.5 | -0.6 to -3.5 | -0.6 to -3.5 | -0.6 to -3.5 | -0.6 to -3.5 | -0.6 to -3.5 | -0.6 to -3.5 |
| Pixel size (Å) | 0.685 | 0.685 | 0.685 | 0.685 | 0.685 | 0.685 | 0.685 | 0.685 |
| Symmetry imposed | C1 | C1 | C1 | C1 | C1 | C1 | C1 | C1 |
| Initial particle images (no.) | 3,254,352 | 3,254,352 | 3,254,352 | 3,254,352 | 3,254,352 | 3,254,352 | 3,254,352 | 3,254,352 |
| Final particle images (no.) | 4,431 | 7,433 | 1,828 | 2,008 | 8,375 | 9,553 | 30,710 | 7,590 |
| Map resolution (Å) | 4.5 | 4.5 | 7.1 | 7.2 | 4.2 | 4.3 | 4.0 | 4.4 |
| FSC threshold | 0.143 | 0.143 | 0.143 | 0.143 | 0.143 | 0.143 | 0.143 | 0.143 |
| Map resolution range (Å) | 4.2-8.0 | 4.2-8.0 | 6.8-15.0 | 7.0-15.0 | 3.5-8.0 | 4.0-8.0 | 3.8-8.0 | 4.2-8.0 |
| **Refinement** | | | | | | | | |
| Initial model used | | | | | | | | |
| Model resolution (Å) | | | | | | | 4.3 | |
| FSC threshold | | | | | | | 0.5 | |
| Model resolution range (Å) | | | | | | | 3.8-8.0 | |
| Map sharpening *B* factor (Å$^2$) | 0 | 0 | 0 | 0 | 0 | 0 | 0 | 0 |
| **Model composition** | | | | | | | | |
| Non-hydrogen atoms | 163221 | 163587 | 163981 | 163421 | 163667 | 163640 | 163694 | 163900 |
| Protein residues | 20671 | 20715 | 20766 | 20702 | 20716 | 20716 | 20717 | 20758 |
| Ligands | 24 | 21 | 21 | 20 | 23 | 22 | 23 | 18 |
| *B* factors (Å$^2$) | | | | | | | | |
| Protein | 124.70 | 130.77 | 138.25 | 182.54 | 135.30 | 139.42 | 150.05 | 132.47 |
| Ligand | 125.44 | 142.00 | 156.26 | 171.59 | 176.71 | 167.61 | 151.12 | 118.41 |
| **R.m.s. deviations** | | | | | | | | |
| Bond lengths (Å) | 0.005 | 0.004 | 0.005 | 0.004 | 0.006 | 0.004 | 0.005 | 0.004 |
| Bond angles (°) | 1.135 | 1.112 | 1.111 | 1.104 | 1.176 | 1.127 | 1.113 | 1.115 |
| **Validation** | | | | | | | | |
| MolProbity score | 1.79 | 1.67 | 1.87 | 1.75 | 1.84 | 1.77 | 1.74 | 1.71 |
| Clashscore | 6.57 | 5.21 | 8.27 | 6.44 | 7.07 | 6.15 | 5.81 | 5.59 |
| Poor rotamers (%) | 0.34 | 0.27 | 0.36 | 0.21 | 0.47 | 0.46 | 0.37 | 0.26 |
| **Ramachandran plot** | | | | | | | | |
| Favored (%) | 93.54 | 94.26 | 93.71 | 94.18 | 92.87 | 93.28 | 93.56 | 93.98 |
| Allowed (%) | 6.21 | 5.53 | 6.13 | 5.66 | 6.90 | 6.41 | 6.23 | 5.83 |
| Disallowed (%) | 0.25 | 0.21 | 0.16 | 0.16 | 0.23 | 0.32 | 0.21 | 0.19 |

### Cryo-EM data collection, refinement and validation statistics (continued)

| | $E_B$-$E_{C1}$ (EMD-62106) (PDB 9K5W) | $E_B$-$E_{C2}$ (EMD-62107) (PDB 9K5X) | $E_B$-$E_{D0}$ (EMD-62108) (PDB 9K5Y) | $E_B$-$E_{D1}$ (EMD-62109) (PDB 9K5Z) | $E_B$-$E_{D2}$ (EMD-62110) (PDB 9K60) | $E_{C1}$-$E_{C1}$ (EMD-62111) (PDB 9K61) | $E_{C1}$-$E_{C2}$ (EMD-62112) (PDB 9K62) | $E_{C1}$-$E_{D0}$ (EMD-62113) (PDB 9K63) |
|---|---|---|---|---|---|---|---|---|
| **Data collection and processing** | | | | | | | | |
| Magnification | 105,000 | 105,000 | 105,000 | 105,000 | 105,000 | 105,000 | 105,000 | 105,000 |
| Voltage (kV) | 300 | 300 | 300 | 300 | 300 | 300 | 300 | 300 |
| Electron exposure (e–/Å$^2$) | 44 | 44 | 44 | 44 | 44 | 44 | 44 | 44 |
| Defocus range (μm) | -0.6 to -3.5 | -0.6 to -3.5 | -0.6 to -3.5 | -0.6 to -3.5 | -0.6 to -3.5 | -0.6 to -3.5 | -0.6 to -3.5 | -0.6 to -3.5 |
| Pixel size (Å) | 0.685 | 0.685 | 0.685 | 0.685 | 0.685 | 0.685 | 0.685 | 0.685 |
| Symmetry imposed | C1 | C1 | C1 | C1 | C1 | C1 | C1 | C1 |
| Initial particle images (no.) | 3,254,352 | 3,254,352 | 3,254,352 | 3,254,352 | 3,254,352 | 3,254,352 | 3,254,352 | 3,254,352 |
| Final particle images (no.) | 1,440 | 1,882 | 9,807 | 12,032 | 44,937 | 1,465 | 2,257 | 7,543 |
| Map resolution (Å) | 7.5 | 7.3 | 4.3 | 4.3 | 4.0 | 9.0 | 6.6 | 4.3 |
| FSC threshold | 0.143 | 0.143 | 0.143 | 0.143 | 0.143 | 0.143 | 0.143 | 0.143 |
| Map resolution range (Å) | 7.0-15.0 | 7.0-15.0 | 4.0-8.0 | 4.0-8.0 | 3.8-8.0 | 8.8-15.0 | 6.0-15.0 | 4.1-8.0 |
| **Refinement** | | | | | | | | |
| Initial model used | | | | | | | | |
| Model resolution (Å) | | | | | 4.3 | | | |
| FSC threshold | | | | | 0.5 | | | |
| Model resolution range (Å) | | | | | 3.8-8.0 | | | |
| Map sharpening *B* factor (Å$^2$) | 0 | 0 | 0 | 0 | 0 | 0 | 0 | 0 |
| **Model composition** | | | | | | | | |
| Non-hydrogen atoms | 164350 | 163793 | 164033 | 164006 | 164007 | 164794 | 164237 | 164480 |
| Protein residues | 20810 | 20746 | 20760 | 20760 | 20760 | 20862 | 20798 | 20812 |
| Ligands | 18 | 17 | 20 | 19 | 20 | 18 | 17 | 20 |
| *B* factors (Å$^2$) | | | | | | | | |
| Protein | 178.93 | 182.77 | 149.48 | 145.54 | 155.64 | 354.19 | 154.30 | 144.85 |
| Ligand | 169.33 | 185.49 | 186.92 | 162.19 | 161.51 | 443.35 | 148.59 | 198.15 |
| **R.m.s. deviations** | | | | | | | | |
| Bond lengths (Å) | 0.004 | 0.004 | 0.005 | 0.005 | 0.005 | 0.005 | 0.004 | 0.004 |
| Bond angles (°) | 1.116 | 1.111 | 1.135 | 1.156 | 1.114 | 1.116 | 1.106 | 1.102 |
| **Validation** | | | | | | | | |
| MolProbity score | 1.84 | 1.87 | 1.80 | 1.83 | 1.73 | 1.96 | 1.78 | 1.67 |
| Clashscore | 7.75 | 8.24 | 6.53 | 6.93 | 5.57 | 9.67 | 6.83 | 4.95 |
| Poor rotamers (%) | 0.31 | 0.35 | 0.38 | 0.43 | 0.32 | 0.26 | 0.29 | 0.30 |
| **Ramachandran plot** | | | | | | | | |
| Favored (%) | 93.68 | 93.54 | 93.29 | 92.95 | 93.46 | 93.03 | 94.00 | 93.78 |
| Allowed (%) | 6.17 | 6.26 | 6.50 | 6.77 | 6.32 | 6.78 | 5.84 | 5.98 |
| Disallowed (%) | 0.15 | 0.20 | 0.21 | 0.29 | 0.21 | 0.19 | 0.16 | 0.24 |

### Cryo-EM data collection, refinement and validation statistics (continued)

| | $E_{C1}$-$E_{D1}$ (EMD-62114) (PDB 9K64) | $E_{C1}$-$E_{D2}$ (EMD-62115) (PDB 9K65) | $E_{C2}$-$E_{C2}$ (EMD-62116) (PDB 9K66) | $E_{C2}$-$E_{D0}$ (EMD-62117) (PDB 9K67) | $E_{C2}$-$E_{D1}$ (EMD-62118) (PDB 9K68) | $E_{C2}$-$E_{D2}$ (EMD-62119) (PDB 9K69) | $E_{D0}$-$E_{D0}$ (EMD-62120) (PDB 9K6A) | $E_{D0}$-$E_{D1}$ (EMD-62121) (PDB 9K6B) |
|---|---|---|---|---|---|---|---|---|
| **Data collection and processing** | | | | | | | | |
| Magnification | 105,000 | 105,000 | 105,000 | 105,000 | 105,000 | 105,000 | 105,000 | 105,000 |
| Voltage (kV) | 300 | 300 | 300 | 300 | 300 | 300 | 300 | 300 |
| Electron exposure (e–/Å$^2$) | 44 | 44 | 44 | 44 | 44 | 44 | 44 | 44 |
| Defocus range (μm) | -0.6 to -3.5 | -0.6 to -3.5 | -0.6 to -3.5 | -0.6 to -3.5 | -0.6 to -3.5 | -0.6 to -3.5 | -0.6 to -3.5 | -0.6 to -3.5 |
| Pixel size (Å) | 0.685 | 0.685 | 0.685 | 0.685 | 0.685 | 0.685 | 0.685 | 0.685 |
| Symmetry imposed | C1 | C1 | C1 | C1 | C1 | C1 | C1 | C1 |
| Initial particle images (no.) | 3,254,352 | 3,254,352 | 3,254,352 | 3,254,352 | 3,254,352 | 3,254,352 | 3,254,352 | 3,254,352 |
| Final particle images (no.) | 12,175 | 10,697 | 1,312 | 7,358 | 9,829 | 11,377 | 14,616 | 43,233 |
| Map resolution (Å) | 4.3 | 4.4 | 8.0 | 4.3 | 4.3 | 4.4 | 4.2 | 3.9 |
| FSC threshold | 0.143 | 0.143 | 0.143 | 0.143 | 0.143 | 0.143 | 0.143 | 0.143 |
| Map resolution range (Å) | 4.0-8.0 | 4.2-8.0 | 7.0-15.0 | 4.0-8.0 | 4.0-8.0 | 4.2-8.0 | 3.8-8.0 | 3.5-8.0 |
| **Refinement** | | | | | | | | |
| Initial model used | | | | | | | | |
| Model resolution (Å) | | | | | | | 4.3 | 4.1 |
| FSC threshold | | | | | | | 0.5 | 0.5 |
| Model resolution range (Å) | | | | | | | 3.8-8.0 | 3.6-8.0 |
| Map sharpening *B* factor (Å$^2$) | 0 | 0 | 0 | 0 | 0 | 0 | 0 | 0 |
| **Model composition** | | | | | | | | |
| Non-hydrogen atoms | 164450 | 164451 | 163680 | 163920 | 163893 | 163894 | 164166 | 164139 |
| Protein residues | 20812 | 20812 | 20734 | 20748 | 20748 | 20748 | 20762 | 20762 |
| Ligands | 19 | 20 | 16 | 19 | 18 | 19 | 22 | 21 |
| *B* factors (Å$^2$) | | | | | | | | |
| Protein | 149.79 | 140.10 | 262.00 | 171.52 | 142.60 | 155.64 | 541.60 | 155.96 |
| Ligand | 170.60 | 156.15 | 218.44 | 157.35 | 168.14 | 159.28 | 737.99 | 177.80 |
| **R.m.s. deviations** | | | | | | | | |
| Bond lengths (Å) | 0.004 | 0.004 | 0.004 | 0.005 | 0.004 | 0.005 | 0.004 | 0.005 |
| Bond angles (°) | 1.126 | 1.114 | 1.110 | 1.161 | 1.110 | 1.157 | 1.115 | 1.133 |
| **Validation** | | | | | | | | |
| MolProbity score | 1.75 | 1.75 | 1.88 | 1.84 | 1.72 | 1.85 | 1.74 | 1.80 |
| Clashscore | 5.63 | 5.61 | 8.52 | 7.12 | 5.55 | 7.08 | 5.70 | 6.31 |
| Poor rotamers (%) | 0.35 | 0.29 | 0.24 | 0.39 | 0.34 | 0.39 | 0.43 | 0.46 |
| **Ramachandran plot** | | | | | | | | |
| Favored (%) | 93.15 | 93.17 | 93.72 | 92.97 | 93.65 | 92.59 | 93.45 | 92.98 |
| Allowed (%) | 6.64 | 6.59 | 6.08 | 6.76 | 6.11 | 7.21 | 6.28 | 6.69 |
| Disallowed (%) | 0.21 | 0.23 | 0.20 | 0.27 | 0.24 | 0.21 | 0.26 | 0.33 |

### Cryo-EM data collection, refinement and validation statistics (continued)

| | $E_{D0}$-$E_{D2}$ (EMD-62122) (PDB 9K6C) | $E_{D1}$-$E_{D1}$ (EMD-62123) (PDB 9K6D) | $E_{D1}$-$E_{D2}$ (EMD-62124) (PDB 9K6E) | $E_{D2}$-$E_{D2}$ (EMD-62125) (PDB 9K6F) |
|---|---|---|---|---|
| **Data collection and processing** | | | | |
| Magnification | 105,000 | 105,000 | 105,000 | 105,000 |
| Voltage (kV) | 300 | 300 | 300 | 300 |
| Electron exposure (e–/Å$^2$) | 44 | 44 | 44 | 44 |
| Defocus range (μm) | -0.6 to -3.5 | -0.6 to -3.5 | -0.6 to -3.5 | -0.6 to -3.5 |
| Pixel size (Å) | 0.685 | 0.685 | 0.685 | 0.685 |
| Symmetry imposed | C1 | C1 | C1 | C1 |
| Initial particle images (no.) | 3,254,352 | 3,254,352 | 3,254,352 | 3,254,352 |
| Final particle images (no.) | 53,317 | 33,571 | 72,897 | 106,179 |
| Map resolution (Å) | 3.9 | 4.0 | 3.9 | 4.0 |
| FSC threshold | 0.143 | 0.143 | 0.143 | 0.143 |
| Map resolution range (Å) | 3.5-8.0 | 3.5-8.0 | 3.5-8.0 | 3.5-8.0 |
| **Refinement** | | | | |
| Initial model used | | | | |
| Model resolution (Å) | 4.2 | 4.3 | 4.1 | 4.2 |
| FSC threshold | 0.5 | 0.5 | 0.5 | 0.5 |
| Model resolution range (Å) | 3.7-8.0 | 3.8-8.0 | 3.6-8.0 | 3.7-8.0 |
| Map sharpening *B* factor (Å$^2$) | 0 | 0 | 0 | 0 |
| **Model composition** | | | | |
| Non-hydrogen atoms | 164140 | 164112 | 164113 | 164114 |
| Protein residues | 20762 | 20762 | 20762 | 20762 |
| Ligands | 22 | 20 | 21 | 22 |
| *B* factors (Å$^2$) | | | | |
| Protein | 157.68 | 141.02 | 153.12 | 167.04 |
| Ligand | 175.01 | 156.21 | 150.41 | 163.87 |
| **R.m.s. deviations** | | | | |
| Bond lengths (Å) | 0.005 | 0.005 | 0.006 | 0.005 |
| Bond angles (°) | 1.126 | 1.149 | 1.155 | 1.127 |
| **Validation** | | | | |
| MolProbity score | 1.78 | 1.85 | 1.84 | 1.78 |
| Clashscore | 6.16 | 7.03 | 6.68 | 5.98 |
| Poor rotamers (%) | 0.44 | 0.60 | 0.52 | 0.43 |
| **Ramachandran plot** | | | | |
| Favored (%) | 93.08 | 92.67 | 92.34 | 92.79 |
| Allowed (%) | 6.68 | 6.91 | 7.29 | 6.95 |
| Disallowed (%) | 0.25 | 0.42 | 0.37 | 0.26 |

**Movie S1.**

Motion illustration of the ubiquitin-proteasome interactions at subunits RPN1, RPN2, RPN10 and α5.

**Movie S2.**

Motion illustration of the structural features of single-nucleotide exchange during translocation initiation.

**Movie S3.**

Motion illustration of the conformational changes of the 26S proteasome during substrate translocation by connecting all states resolved in this study.
